# Paradoxical energetics in the polar diatom *Fragilariopsis cylindrus* exposed to extreme low light

**DOI:** 10.64898/2026.01.30.702734

**Authors:** Arthur Plassart, Nathalie Joli, Sneha Sivaram, Sébastien Guérin, Flavienne Bruyant, Marie-Hélène Forget, Chris Bowler, Marcel Babin

## Abstract

In the Arctic Ocean, diatoms initiate blooms under sea ice at extreme low-light levels, yet the limits and mechanisms behind this capability remain unknown. We investigated the steady-state physiological and molecular responses of the polar diatom *Fragilariopsis cylindrus* across a light gradient (0.1 to 30 µmol photons m^-2^ s^-1^), representative of under-ice winter to early spring conditions, and reveals distinct strategies to cope with low light at both ends of this range. While cells optimize photon capture efficiency between 3 and 15 µmol photons m^-2^ s^-1^ relative to 30 µmol photons m^-2^ s^-1^, this strategy collapses below 1 µmol photons m^-2^ s^-1^. In this dim-light regime, cells activate non-photochemical quenching and a sustained xanthophyll cycle, which indicates a paradoxical requirement for energy dissipation despite extreme photon scarcity. While cell division arrests at 0.18 µmol photons m^-2^ s^-1^, photosynthetic electron transport seems to remain possible down to 0.1 µmol photons m^-2^ s^-1^, which suggests an uncoupling between photosynthesis and biomass accumulation. Crucially, this low-light regime occurs without the consumption of reserves and represents a physiological state distinct from metabolic hypometabolism in prolonged darkness. We propose that this dim-light physiological state arises when residual light absorption exceeds the energetic requirements for cellular maintenance in the absence of division. The result is a regulated imbalance dissipated through heat and carbon excretion. This mechanism allows polar diatoms to maintain a primed photosynthetic metabolism to facilitate rapid growth recovery upon the return of light after the winter solstice.

## INTRODUCTION

Oceanic primary production is fundamentally constrained by the depth where gross primary production balances respiration and thus enables net biomass accumulation. Historically, this limit has been approximated by the euphotic zone, defined as the depth where light reaches 1% of surface photosynthetically active radiation (PAR) (Behrenfeld & Boss, 2018; Sverdrup, 1953). However, this relative threshold is physiologically arbitrary and exceeds the minimal light requirements for many phototrophs (Marra et al., 2023; Banse, 2004). Indeed, in the Arctic, recent field observations confirm net phototrophic accumulation under sea ice at irradiances far below conventional limits, with division rates sustained between 0.04 and 0.8 µmol photons m^-2^ s^-1^ (Hoppe et al., 2024; Randelhoff et al., 2020; Kvervink et al., 2018; Hancke et al., 2018). These findings, reveal that polar phytoplankton reinitiate growth in near-darkness and thus imply that the theoretical minimum quantum of light requirement for oxygenic photosynthesis, modeled to be at 0.01 µmol photons m^-2^ s^-1^ (Raven et al., 2000), is reached very early in the season, under the ice during the polar winter. In this context, diatoms are the dominant contributors to the sympagic (ice-associated) and pelagic phytoplankton communities and play a central role in polar ecosystems through both their abundance and productivity (Ardyna and Arrigo, 2020; Lafond et al., 2019; Leu et al., 2015; Alou Font et al., 2013; Poulin et al., 2011; Lovejoy et al., 2002). Despite their ecological importance, the physiological limits of photosynthetic growth at extremely low irradiance of light remain unknown for these, and other, unicellular photoautotrophs found in Arctic ecosystems.

Existing paradigms of photoacclimation derive largely from studies at moderate low-light irradiance (typically ∼5-15 µmol photons m^-2^ s^-1^). In these regimes, unicellular algae such as diatoms maximize photon capture by increasing the cellular content of photosynthetic pigments, thereby improving photosynthetic efficiency (Agarwal et al., 2023; Croteau et al., 2021; Hasley et al., 2015; Arrigo et al., 2010; Suggett et al., 2007; Quigg et al., 2006; Dubinsky et al., 1986; Geider et al., 1985, 1986; Laws & Bannister, 1980; Falkowski and Owens, 1980). Furthermore, they simultaneously downregulate respiratory costs to lower the compensation point for growth (Hasley et al., 2015; Dubinsky & Stambler, 2009; Quigg & Beardall, 2003; Fisher et al., 1989). In sharp contrast, the response to complete darkness involves a fundamental shift toward hypometabolism, which relies on the consumption of endogenous reserves. Under these conditions, specifically during the polar night, Arctic diatoms arrest their cell cycle, degrade photosynthetic machinery, and actively catabolize intracellular reserves (such as lipids and proteins) via *β*-oxidation and autophagy to sustain essential housekeeping functions (Joli et al., 2024; Juchem et al., 2023; Kennedy et al., 2019; Sciandra et al., 2022; Morin et al., 2020; Lacour et al., 2019; Schaub et al., 2017). Between these two extremes in light conditions (darkness, and low to high light), very low light remains a regime where photosynthesis is largely unexplored, yet it is highly relevant to the under-ice polar environment. Specifically, it remains unclear whether cells maintain phototrophic growth in a dim-light regime (below 5 µmol photons m^-2^ s^-1^) or if they perceive scarce photons as total darkness and thus trigger the dark-acclimation processes described above. As Arctic ice cover declines in extent, thickness and seasonal persistence due to climate change, light conditions will change significantly in the Arctic Ocean (Kristiansen et al., 2025; Castellani et al., 2022; Lewis et al., 2020). Consequently, an understanding of the physiological limits and responses of these key diatom producers is critical to predict future polar ecosystem dynamics.

To determine the absolute lower limit of photosynthetic growth, we investigate the steady-state physiological and molecular responses of the polar diatom *Fragilariopsis cylindrus* after full acclimation to a range of low light intensities. These intensities range from an extreme “dim-light” regime (0.1-1 µmol photons m^-2^ s^-1^), to low light (3-15 µmol photons m^-2^ s^-1^) and a moderate condition (30 µmol photons m^-2^ s^-1^), representative of under-ice light levels between the winter solstice and early spring conditions (Cohen et al., 2020; Massicotte et al., 2020). Through the integration of growth kinetics, photophysiology, and metabolic profiles, we designed our experiments specifically to capture the physiological threshold that separates light-limited growth from growth arrest. Our results reveal two divergent acclimation strategies. While cells optimize photon capture efficiency under low light, this classical photoacclimation response collapses in the dim-light regime. Indeed, under these conditions, cells neither divide nor enhance light harvest. Rather, they activate non-photochemical quenching (NPQ) and a sustained xanthophyll cycle. In the context of the extreme Arctic light cycle, this distinct dim-light strategy may be central to the ecological success of polar diatoms, allowing them to resume growth immediately once light levels exceed the compensation threshold and thereby preventing the lag phases associated with recovery from deep hypometabolism.

## RESULTS

### Cellular growth and physiology

*F. cylindrus* cultures were beforehand acclimated to six continuous light levels ranging from 0.1 to 30 µmol photons m^-2^ s^-1^ for 20 days. During this period, we measured cell density, chlorophyll content and basic photophysiology parameters. The growth rate (μ) followed a typical logarithmic light-dependent response, peaking at a maximum rate (μ_max_) of 0.10 ± 0.01 d^-1^ at 30 µmol photons m^-2^ s^-1^ (Figure S1, Table S2). The light saturation parameter for growth (𝐸^*K*^_μ_), which marks the transition between light-limited and saturating conditions, was estimated at 4.71 ± 1.74 µmol photons m^-2^ s^-1^ (Figure S1, Table S2). This 𝐸^*K*^_μ_ value is low but typical of values reported for polar taxa (Lacour et al., 2017). The monitoring of cell density indicates that while cultures at 0.3 µmol photons m^-2^ s^-1^ maintained a modest but positive growth, cell division ceased at 0.1 µmol photons m^-2^ s^-1^ (Figure 1A, Table S1). We estimated, using a mathematical model, a light compensation point for growth (I_c_) of 0.18 ± 0.11 µmol photons m^-2^ s^-1^, this theoretical threshold aligns closely with the observed experimental arrest of cell division (Figure 1A). Despite the arrest of growth at the lowest irradiances, cell viability remained high (>90%) across all treatments (Figure 1B), with only minor declines at 0.1, 0.3, and 30 µmol photons m^-2^ s^-1^. Furthermore, since the dark-adapted quantum yield for photochemistry in PSII (F_v_/F_m_) did not show a drastic drop with decreasing light, these cells did not appear to be in a significant physiological stress (Figure 1C). Morphologically, cell size significantly decreased at low-light irradiances (3-15 µmol photons m^-2^ s^-1^) relative to with high light conditions, but increased again under dim-light conditions (<1 µmol photons m^-2^ s^-1^) to reach values comparable to the high-light conditions (Figure 1D, Figure S2A). Transcriptional profiles show that the machinery required for DNA replication, such as the origin recognition complex (ORC) and the proliferating cell nuclear antigen (PCNA), was maintained even at the growth-arrest irradiance of 0.1 µmol photons m^-2^ s^-1^. Similarly, transcripts for general cell cycle regulators, including the Skp1/Cullin/F-box complex (SCF), structural maintenance of chromosomes (SMC) proteins, and the Anaphase-Promoting Complex core subunits (APC/C) with its co-activator CDH1, were also stable (Figure S4A). In contrast, the transcript level of CDC20 genes was almost undetectable, which suggests that the specific absence of this component may contribute to the arrest of cell division in dim-light conditions (Figure S4A). Interestingly a similar observation was made in cells of *F. cylindrus* acclimated to prolonged darkness (Figure S5B), where cells entered in a quiescent state without cell division.

**Figure 1.**
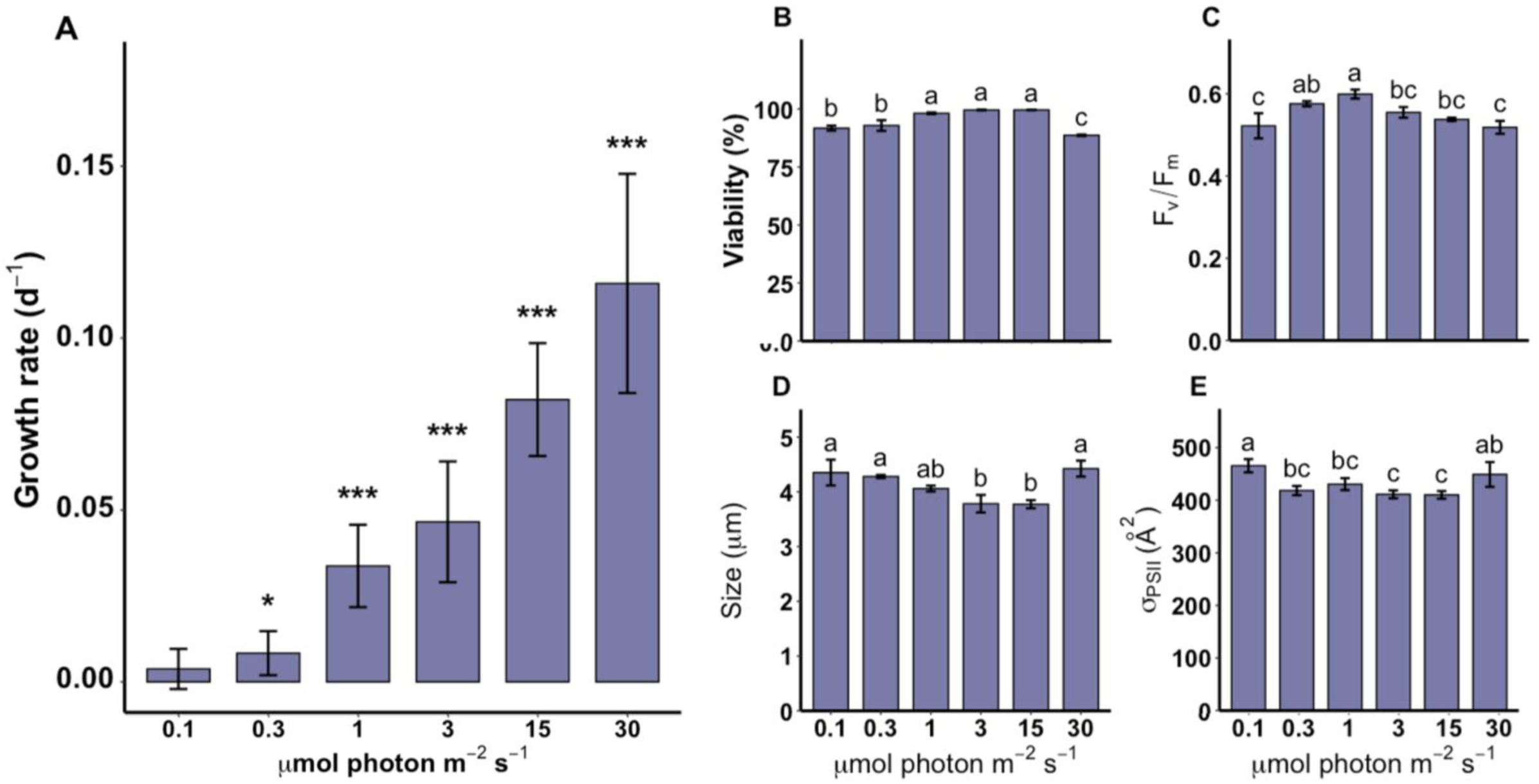
Influence of irradiance on growth dynamics and physiological traits. (A) Specific growth rate (µ, d^-1^). (B) Percentage of viable cells (%). (C) Maximum quantum yield of photosystem II (F_v_/F_m_). (D) Cell size (μm). (E) Functional absorption cross-section of PSII (σPSII, in Å^2^). Blue bars represent means (n=3). In (A), error bars represent 95% confidence intervals and asterisks indicate growth rates significantly different from zero (p-value of t-test, ∗<0.05 & ∗∗∗<0.001). In (B-E), error bars represent standard deviation (SD) and different letters indicate significant differences between irradiance levels (P<0.05) determined by Tukey’s HSD or Dunn’s post hoc analysis.

### Bioenergetics and central metabolism

Biovolume-normalized quotas of organic carbon (*Q^C^_V_*) and nitrogen (*Q^N^_V_*) remained stable across the light gradient (Figure 2A, B), which indicates that cells maintained stable internal C and N organic pools even at growth-arrest irradiance. This observation argues against an endogenous catabolism shift driven by the degradation of organic reserves to sustain basal metabolism, a strategy previously reported in *F. cylindrus* after exposure to prolonged darkness (Joli et al., 2024; Kennedy et al., 2019; Sciandra et al., 2022; Morin et al., 2020). Consistently, transcriptomic profiles indicated that the core metabolic architecture remained remarkably conserved across all light conditions. The number of expressed genes (counts > 30) within major metabolic pathways was largely identical along the light gradient (Figure 4). However, a generalized decrease in transcript abundance was observed across pathways (Z-score value in Figure 4; Figure S6), consistent with a global metabolic slowdown rather than a qualitative reorganization of cellular metabolism. Notably, genes associated with the Calvin-Benson-Bassham (CBB) cycle, lipid biosynthesis, and mitochondrial *β*-oxidation remained actively transcribed even under the lowest light conditions (Figure 4; Figure S7). This suggests that central anabolic and catabolic pathways remain functional despite growth arrest in the dimmest light regime. Microscopy highlighted a distinct phenotypic shift at low light, characterized by increased cell aggregation (Figure S2B-C). Accordingly, genes within Extracellular Polymeric Substance (EPS) biosynthesis pathways were upregulated in growth-arrested cells (Figure 4; Figure 5). This response aligns with stress-related strategies commonly reported in sea ice diatoms under extreme environmental constraints (Aslam et al., 2017; Aslam et al., 2012; Underwood et al., 2010). Taken together, these results indicate that *F. cylindrus* does not undergo a major metabolic reconfiguration under dim light but instead maintains its core metabolic machinery in a low-activity, homeostatic state.

**Figure 2.**
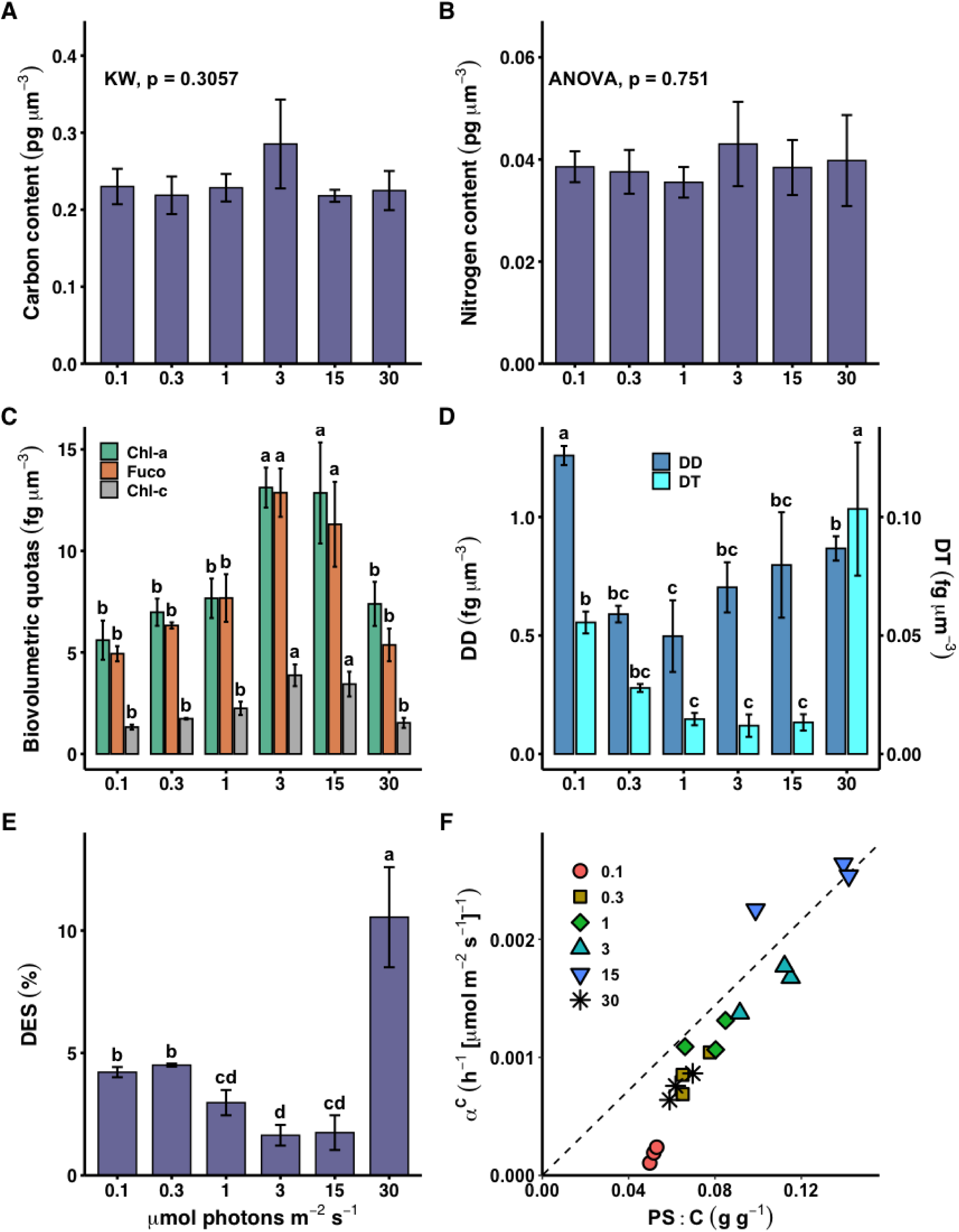
Influence of growth irradiance on biovolumetric quotas and pigment stoichiometry. (A & B) Cellular organic carbon and nitrogen biovolumetric quotas. (C) Biovolumetric quotas of photosynthetic pigments: chlorophyll *a* (Chl *a*, green), fucoxanthin (Fuco, orange), and chlorophyll c_1_ + c_2_ (Chl *c*, grey). (D) Biovolumetric quotas of photoprotective pigments: diadinoxanthin (DD, dark blue) and diatoxanthin (DT, cyan). (E) De-epoxidation state (DES), calculated as [DT / (DD + DT)] × 100. (F) Relationship between the carbon-specific initial slope of the photosynthesis versus irradiance curve (α^C^) and the total photosynthetic pigment to carbon ratio (PS:C). Symbols represent distinct growth irradiances (biological triplicates). In C-E, bars represent means ± SD (n = 3), and different letters indicate significant differences between irradiance levels (P < 0.05) determined by Tukey’s HSD or Dunn’s post hoc analysis.

**Figure 3.**
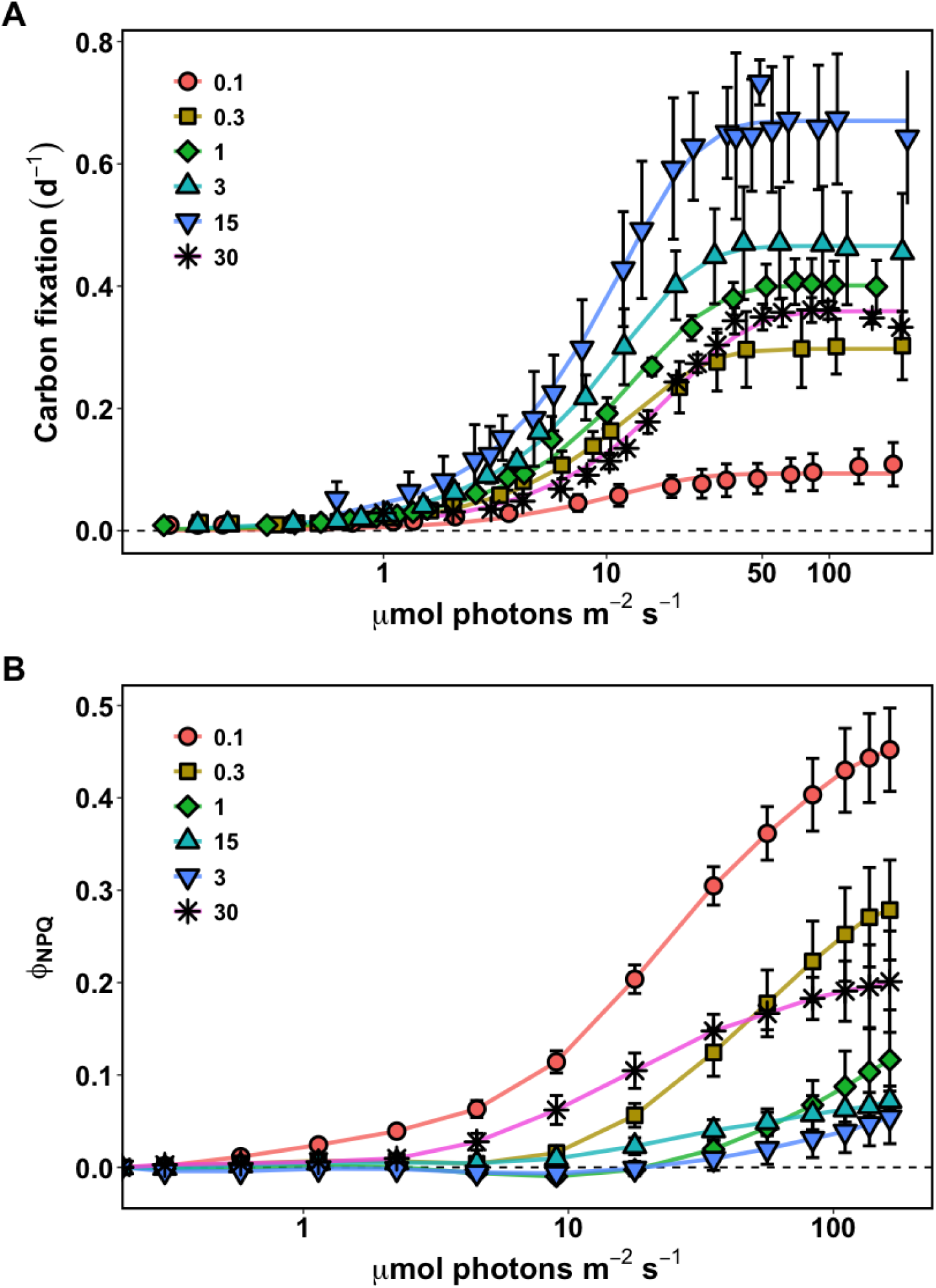
Influence of growth irradiance on photosynthesis and light utilization. (A) ^14^C uptake versus incubation irradiance curves. Symbols represent mean fixation rates normalized by cellular organic carbon content for cells acclimated to 0.1, 0.3, 1, 3, 15, and 30 μ mol photons m^-2^ s^-1^. Curves represent the modelled carbon fixation (Jassby and Platt, 1976). (B) Non-photochemical quenching yield (ϕNPQ) during rapid light curves, derived as ϕNPQ=F′/Fm′−F′/FM. Data represent means ± SD (n=9), and lines connect mean values.

**Figure 4.**
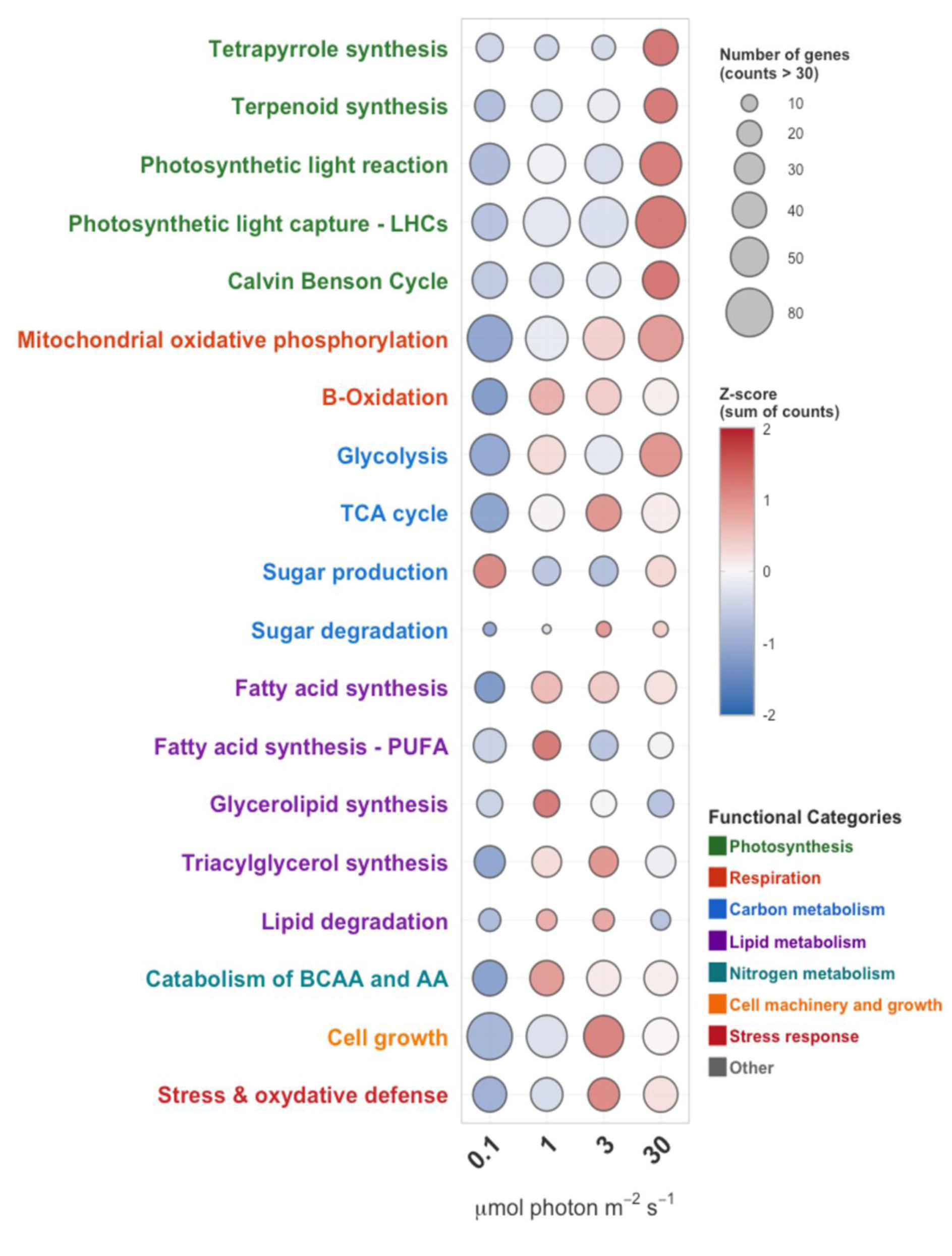
Transcriptional patterns of metabolic pathways following the acclimation period specific to each growth irradiance. The bubble plot summarizes the expression profiles of 898 genes grouped by metabolic pathway (see Dataset S1 for the detailed gene list). Analysis was performed on DESeq2-normalized counts. Pathway text labels are color-coded by broad functional category: Photosynthesis (green), Respiration (red), Carbon metabolism (blue), Lipid metabolism (violet), Amino acid metabolism (cyan), Cell machinery and growth (orange), and Stress response (dark red). Bubble size is proportional to the number of expressed genes (>30 normalized counts) detected within the pathway. The bubble colour gradient represents the relative expression level (Z-score) of the cumulative pathway abundance. The Z-score is calculated as Z=(X−μ)/σ, where X is the sum of normalized counts for a pathway at a specific irradiance, and μ and σ represent the mean and standard deviation of these sums across all treatments, respectively.

**Figure 5.**
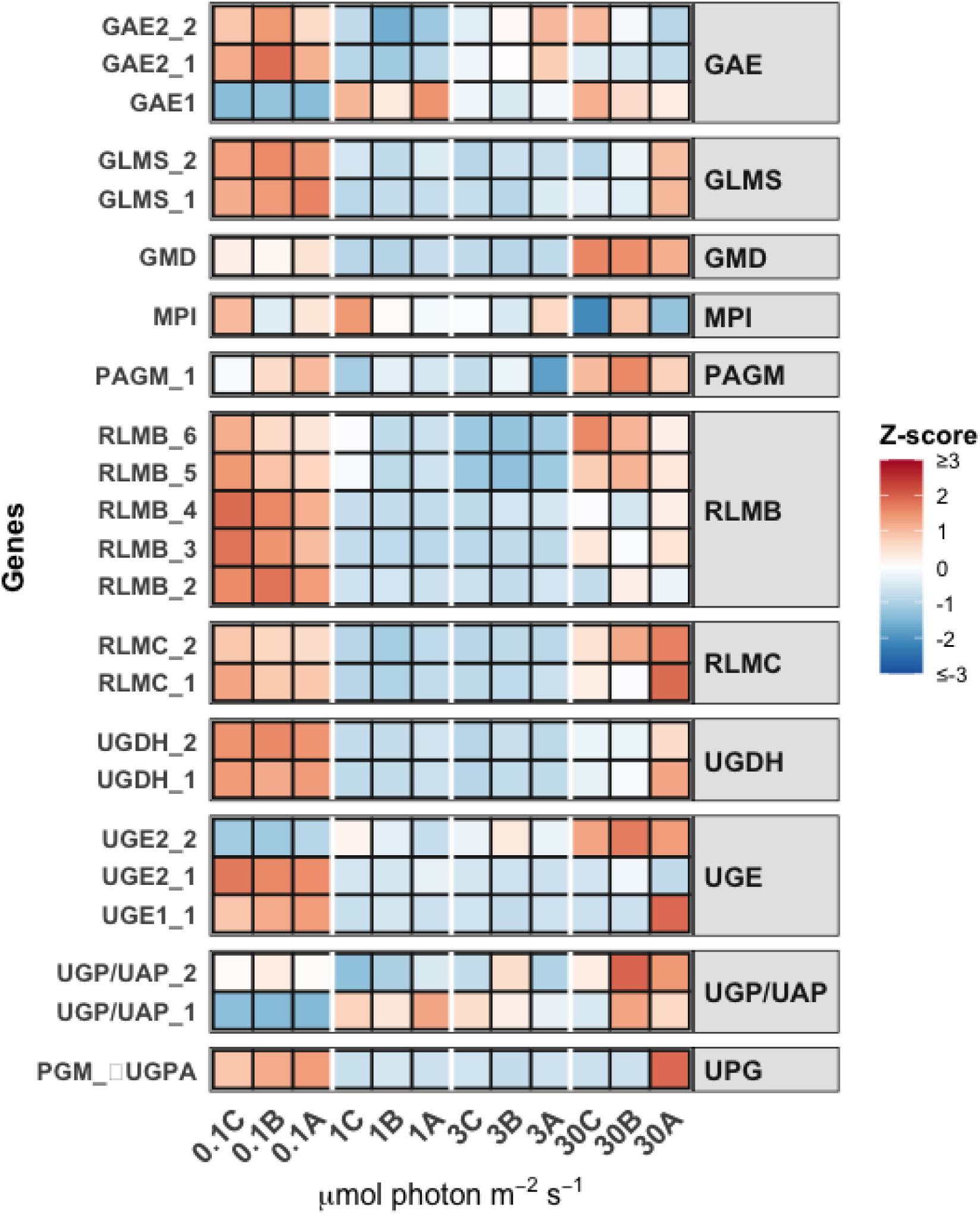
Transcriptional profiling of genes involved in EPS biosynthesis pathways. The heatmap shows the expression patterns of genes involved in exopolysaccharide (EPS) biosynthesis in *F. cylindrus*, identified according to Aslam et al. (2017), following the acclimation period specific to each growth irradiance. The colour gradient indicates the relative expression level (Z-score) for each gene across all biological triplicates and light conditions. Abbreviations: GAE, UDP-glucuronate 4-epimerase; GLMS, glutamine-fructose-6-phosphate transaminase; GMD, GDP-mannose 4,6-dehydratase; MPI, mannose-6-phosphate isomerase; PAGM, phosphoacetylglucosamine mutase; RLMB, dTDP-glucose 4,6-dehydratase; RLMC, bifunctional dTDP-4-dehydrorhamnose 3,5-epimerase/TDP-4-dehydrorhamnose reductase; UGDH, UDP-glucose 6-dehydrogenase; UGE, UTP-glucose-1-phosphate uridylyltransferase; UGP/UAP, UDP-N-acetylglucosamine diphosphorylase/UTP-glucose-1-phosphate uridylyltransferase. See dataset S1 for the corresponding JGI ID gene.

### Photosynthetic pigments and photosynthetic performance

Photosynthetic pigments exhibited marked variations in response to changes in light availability. Quotas of chlorophyll *a* (Chl *a*) and accessory pigments (fucoxanthin (Fx) and chlorophyll c_1_ + c_2_) were approximately 2.4-fold higher in low-light acclimated cells (3-15 µmol photons m^-2^ s^-1^) compared to values at 30 µmol photons m^-2^ s^-1^ (Figure 2C), which is a typical low-light acclimation strategy (Agarwal et al., 2023; Arrigo et al., 2010; Kropuenske et al., 2009; Dubinsky & Stambler, 2009; Quigg et al., 2006; Falkowski and Owens, 1980). This accumulation was accompanied by a shift in pigment stoichiometry, with a slight increase in Fx relative to Chl *a* (Figure S3A). At the transcriptional level, this low-light response (3 and 1 µmol photons m^-2^ s^-1^) was associated with the specific induction of genes that encode *Lhcf*25 (ProtID: 193422) and *Lhcf*26 (ProtID: 195777) (Figure S9). These adjustments consistently correlated with the increase in the carbon-specific initial slope (α^C^) of the photosynthesis-irradiance response (PE) curve at 1 to 15 µmol photons m^-2^ s^-1^, a clear sign of optimization in light harvest for photochemistry (Figure 2F). This photoacclimation strategy was, however, not found in dim-light conditions. While pigment quotas at 0.1 µmol photons m^-2^ s^-1^ were similar to those at 30 µmol photons m^-2^ s^-1^ (Figure 2C), the α^C^ coefficient and the light-saturated carbon-specific photosynthetic rate (𝑃^𝐶^) declined dramatically (Figure 2F, Figure 3A, Figure S4). The linear relationship between α^C^ and photosynthetic pigments (PS:C ratio) found at 3-15 µmol photons m^-2^ s^-1^ broke down at 0.1 µmol photons m^-2^ s^-1^ (Figure 2F), with a loss of light use efficiency for photochemistry despite the maintenance of the pigment pool and core machinery transcripts. This physiological collapse contrasts with the transcriptomic stability of the core photosynthetic machinery. The nuclear genes encoding for reaction center cores (PSII and PSI), the oxygen-evolving complex (OEC), PSII repair and assembly mechanisms, and the entire thylakoid electron carriers displayed relatively comparable transcript levels across all light conditions (Figure S10). This suggests that the photosynthetic machinery is maintained but functionally decoupled regarding the maximal fixation capacity observed (𝑃^𝐶^, Figure S4). Additionally, genes for the different types of antennae showed a distinct signature under reduced light, as the majority of *Lhcf* genes (PSII antennae) were downregulated at 0.1 µmol photons m^-2^ s^-1^, while *Lhcr* gene transcripts (PSI antennas) remained significantly more abundant than *Lhcf* transcripts (Figure S9).

### Xanthophyll pigments and NPQ

The loss of photosynthetic efficiency at 0.1 µmol photons m^-2^ s^-1^ coincided with an activation of photoprotective mechanisms. While light-harvest pigment quotas increased with decreasing light intensity and remained stable in the dim-light condition, xanthophyll cycle pigments showed the opposite trend (Figure 2D). The total pool of diadinoxanthin (Dd) and diatoxanthin (Dt) declined by ∼30% as irradiance decreased from 30 to 0.3 µmol photons m^-2^ s^-1^. In contrast, a specific *de novo* synthesis of xanthophylls was observed at 0.1 µmol photons m^-2^ s^-1^ (Figure 2D). While the de-epoxidation state (DES) dropped between 30 and 1 µmol photons m^-2^ s^-1^, it paradoxically increased significantly at the lowest irradiances (0.1-0.3 µmol photons m^-2^ s^-1^), indicating a reactivation of the xanthophyll cycle to dissipate light energy even at these very low light levels (Figure 2D; Figure 3B). This physiological reactivation of the xanthophyll cycle aligns with the transcriptomic results. In pennate diatoms and also in polar specific diatoms, the thermal dissipation of light energy through non-photochemical quenching (NPQ) relies on both xanthophyll pigment content and *Lhcx* proteins (Bailleul et al., 2010; Croteau et al., 2025). The transcriptomic profile revealed that widely expressed *Lhcx* genes (specifically *Lhcx4*, *-7*, *-8*, and *-10*) maintained extremely high expression levels across all conditions, effectively dominating the transcript pool at 0.1 µmol photons m^-2^ s^-1^ by more than an order of magnitude over the diminished light-harvesting genes (Figure S9). Similarly, genes encoding for the xanthophyll cycle enzymes Violaxanthin De-Epoxidase (VDE) and Zeaxanthin Epoxidase (ZEP) also maintained high transcript levels in all conditions (Figure S11A). In diatoms, these enzymes regulate the conversion of Dd into Dt to trigger energy quenching and the reverse reaction, respectively (Giossi et al., 2025a, 2025b). Notably, in complete darkness, *F. cylindrus* lacks expression of genes that encode VDE (Figure S11B and see Joli et al., 2024). This constitutive investment in photoprotective machinery (*Lhcx* and enzymes), combined with the re-accumulation of xanthophylls, appears to influence downstream photochemistry in PSII. Rapid Light Curves (RLC) revealed that, while cells in the photoacclimation window (1-15 µmol photons m^-2^ s^-1^) maintained low induction of NPQ, cells at 0.1 µmol photons m^-2^ s^-1^ exhibited a marked ∼6-fold increase in the quantum yield of non-photochemical quenching (ϕNPQ) compared to cells acclimated to 1-15 µmol photons m^-2^ s^-1^ (Figure 3A, B). This change indicates that the energy absorbed by the maintained antenna complexes was actively diverted from photochemistry (ϕPSII) to regulated heat dissipation upon the cessation of cell division. Furthermore, NPQ induction kinetics confirmed a faster response for non-growing cells, showing NPQ activation at the lowest light levels of the RLC, well before photosynthesis estimated by carbon fixation reached saturation (Figure 3A, B). This strong NPQ engagement provides a mechanistic explanation for the paradoxical loss of photosynthetic efficiency observed for non-dividing cells (Figure 2F).

## DISCUSSION

Through the integration of growth kinetics, photophysiology, and transcriptomic profiles of *F. cylindrus* across a light gradient representative of the sub-ice polar environment, this study defines the physiological thresholds and the photosynthetic and molecular strategies that support diatom survival at the edge of photoautotrophic life. The data suggest that while cells optimize light harvest under low light, they transition to a distinct “dim-light” strategy below the growth compensation point of 0.18 µmol photons m^-2^ s^-1^. In this non-dividing state, *F. cylindrus* paradoxically appears to activate photoprotective quenching based on xanthophyll cycle while the photosynthetic machinery remains in a state of functional readiness. This metabolic homeostasis is apparently distinct from the catabolic hypometabolic state characteristic of the polar night as previously reported (Joli et al., 2024). Our results provide a mechanistic framework for an understanding of how polar diatoms may exploit the earliest Arctic spring photon fluxes, and thus define the limits of photosynthetic growth in extreme environments.

### The limits to growth

The present study identified a growth compensation point (I_c_) of 0.18 ± 0.11 µmol photons.m^-2^ s^-1^ for the polar diatom *F. cylindrus*. This value appears consistent with historical *in situ* observations from Arctic planktonic and sympagic natural assemblages which were based solely on changes in chlorophyll *a* concentrations or carbon acquisition and reported compensation thresholds between ∼0.17 and 0.8 µmol photons m^-2^ s^-1^ (Kvervink et al., 2018; Hancke et al., 2018; Mock & Gradinger, 1999; Palmisano et al., 1985; Suzuki et al., 1997; Cota, 1985). While direct laboratory measurements of light compensation points for polar strains are lacking, a limited number of studies on temperate diatoms report compensation points of similar magnitude, in the range of ∼0.15 to 1 µmol photons m^-2^ s^-1^ (Quigg et al., 2006; Geider et al., 1985; Langdon, 1987, Falkowski and Owens, 1980). This similarity suggests a broadly conserved lower light boundary for growth across diatoms. By contrast, a recent study documented positive biomass accumulation in the water column in the central Arctic basin at a daily irradiance of just 0.04 ± 0.02 µmol photons m^-2^ s^-1^, an order of magnitude below our estimate (Hoppe et al., 2024), although this assessment relied on chlorophyll concentration, an index that may not perfectly reflect changes in biomass. Furthermore, during these periods, the assemblage was dominated by flagellates and dinoflagellates rather than diatoms, which suggests that the observed biomass accumulation might result primarily from these other groups. Whether these taxa possess lower light requirements remains an open question, as, to our knowledge, no light compensation thresholds for growth have yet been reported in the literature for Arctic dinoflagellates, haptophytes, or other flagellate taxa. This highlights a critical knowledge gap in Arctic microalgal ecology. Our empirical threshold remains more than one order of magnitude above the theoretical minimum limit for net photosynthetic growth derived by Raven et al. (2000) of 0.01 µmol photons m^-2^ s^-1^. It should also be noted that, despite the arrest of cell division, the cell population at 0.1 µmol photons m^-2^ s^-1^ exhibited no particular signs of stress or senescence. Ultimately, this extreme low-light growth capability implies that unicellular diatoms in the Arctic could initiate exponential growth very early in the season under sea ice and provides a physiological basis for field observations of active winter growth such as those recently reported (Hoppe et al., 2024; Randelhoff et al., 2020).

### A distinct metabolic state: autotrophy versus dark hypometabolism

The physiological state observed at 0.1 µmol photons m^-2^ s^-1^ appears to differ fundamentally from the hypometabolism response triggered by complete darkness recently reported by Joli et al. (2024). In previous studies, Arctic diatoms in prolonged darkness seem to undergo a switch to cell cycle arrest, the degradation of photosynthetic machinery (proteic actors downstream of the PSII and/or the enzymatic pool involved in the CBB), and the catabolism of intracellular reserves through the *β-*oxidation of lipids and the recycling of proteins and amino acids to sustain housekeeping functions (Joli et al., 2024; Kennedy et al., 2019; Sciandra et al., 2022; Morin et al., 2020). The same trends have been observed in the green microalga *Koliella antartica* and the haptophyte *Phaeocystis antarctica*, and appear to be common to various polar taxa exposed to 3-4 weeks of total darkness (Baldisserotto et al., 2005; van de Pol et al., 2023). In sharp contrast, the current data suggest that cells grown at 0.1 µmol photons m^-2^ s^-1^ maintain stable organic carbon and nitrogen quotas without any apparent signs of chlorosis (Figure 2C). Transcriptomic profiles confirmed that central metabolic pathways remained actively transcribed, albeit at lower abundance (Figure 4). This suggests that *F. cylindrus* does not reconfigure its metabolism toward catabolism but rather maintains a fully functional autotrophic machinery to support basal energy requirements under sub-compensatory irradiance. This metabolic poise aligns with the same principle early observed in the chlorophyte *Dunaliella tertiolecta* (Hellebust and Terborgh, 1967), where low photon fluxes seemed to sustain a reduced but functional protein pool, bypassing the degradative pathways triggered by total darkness. Nevertheless, the fundamental trade-offs in protein allocation required to maintain such a state have remained elusive. In our case, the decrease in 𝑃^𝐶^ for non-dividing cells indicates a drastic decrease in the CBB enzymatic pool (probably Rubisco) and/or protein actors involved in electron transport in the thylakoid photosynthetic chain (Figure 3A). However, the maintenance of a positive light-saturated carbon-specific photosynthetic rate (𝑃_m_^𝐶^) at 0.1 µmol photon m^-2^ s^-1^ supports the presence of a functional protein pool (Figure 3A, Figure S4), which is not the case after one month of full dark acclimation for *F. cylindrus* (Morin et al., 2020). A possible ecological benefit of this strategy is the potential for rapid growth recovery. Previous work on dinoflagellates has demonstrated that populations maintained at sub-compensatory light can recover maximum growth rates almost immediately upon re-illumination, whereas dark-acclimated cells exhibit apparent lag phases (Richardson and Fogg 1982). This active maintenance potentially confers a competitive advantage as it allows for immediate growth recovery once light levels rise above the compensation threshold, and thus prevents the lag phases associated with exiting deep dark hypometabolism.

### Photoprotection at the lower limit of photosynthetic growth: The “Energy Surplus” hypothesis

The most striking feature of this sub-compensatory state represents an apparent paradox in photoacclimation. Typically, low-light acclimation involves the maximization of light harvest and quantum yield (Agarwal et al., 2023; Croteau et al., 2021; Hasley et al., 2015; Arrigo et al., 2010; Suggett et al., 2007; Quigg et al., 2006; Dubinsky et al., 1986; Geider et al., 1985, 1986; Laws & Bannister, 1980; Falkowski and Owens, 1980). However, at 0.1 µmol photons m^-2^ s^-1^, we observed a collapse in both photosynthetic efficiency (α^C^) and photosynthetic maximum capacity (𝑃^𝐶^), which occurs in parallel with a strong upregulation of photoprotective mechanisms, including increased xanthophyll content (Dd+Dt) as well as the sustained expression of *Lhcx* genes. We propose an “Energy Surplus” hypothesis to explain this response. Cellular energy requirements can be partitioned into investment costs (to support cell division, DNA, and organelle duplication) and maintenance costs (which cover protein turnover, ROS detoxification, and homeostasis) (Bender et al., 2022; Raven and Beardall, 2016). Investment in growth represents the primary energy sink for the cell. Under steady-state conditions at 0.1 µmol photons m^-2^ s^-1^, the arrest of cell division apparently eliminates this investment cost. Concurrently, cells appear to minimize maintenance costs through the downregulation of their enzymatic machinery, a classic low-light survival strategy (Hasley et al., 2015; Quigg & Beardall, 2003; Fisher et al., 1989), as evidenced here by the sharp decline in 𝑃^𝐶^, which serves as a proxy for the active Rubisco pool. Furthermore, at 0 °C, the remaining basal metabolic rates are likely to be thermodynamically low. Due to the absence of the primary energy sink (cell division) and the reduction of enzymatic demand, even the minimal electron flow generated at 0.1 µmol photons m^-2^ s^-1^ may exceed the metabolically restricted capacity of the cell to utilize it. To manage this imbalance and help prevent photo-oxidative damage, cells appear to activate specific dissipative sinks. On the one hand, the upregulation of NPQ and xanthophyll synthesis acts as a safety valve for the dissipation of excess excitation energy upstream of the reaction centers, which reflects high-light acclimation responses (Croteau et al., 2026; Croteau et al., 2021; Lacour et al., 2020; Lavaud & Goss, 2014; Lavaud et al., 2002). On the other hand, the upregulation of EPS biosynthesis genes and the observed aggregation (Figure 5, Figure S2) suggest that cells might secrete excess fixed carbon as extracellular polymeric substances, thereby providing a metabolic overflow mechanism, a strategy employed by *F. cylindrus* in the stationary phase (Ugalde et al., 2013).

In conclusion, our study provides a mechanistic definition of the lower light limit of photosynthetic growth in the polar diatom *F. cylindrus*. The results demonstrate that the arrest of cell division at the light compensation point does not trigger metabolic collapse or a shift to dark-induced hypometabolism. Instead, cells enter a poised homeostatic state, where they actively manage relative energy surplus via photoprotective quenching and carbon excretion while they maintain their photosynthetic machinery in functional readiness. These findings suggest that the traditional 1% light threshold is an inappropriate indicator in the Arctic and highlight the requirement for absolute irradiance thresholds to predict the timing of biological activities. However, although the growth limit identified here marks the arrest of cell division at an irradiance still an order of magnitude above the theoretical minimum for maintenance (Raven et al., 2000), photosynthetic activity appears to remain functional. This decoupling between growth and photosynthesis raises a fundamental question about the precise irradiance threshold required for the photosynthetic machinery to truly cease to function. Future research could bridge this gap through the assessment of energy fluxes and acclimation kinetics in *F. cylindrus* and other dominant taxa, such as flagellates and dinoflagellates, during prolonged exposure to sub-compensatory and fluctuating dim light regimes. Through the establishment of the first physiological bricks concerning the ultimate limits of photoautotrophy, this work participates to refine our vision of Arctic microalgal ecology (Hoppe et al., 2024) and provides a framework for the exploration of the potential for photosynthetic life in extreme or extraterrestrial environments (Khuller et al., 2024).

## MATERIALS AND METHODS

### Culture Conditions and Experimental Design

Axenic cultures of the polar diatom *Fragilariopsis cylindrus* (CCMP 3323) were maintained at 0 ± 1°C in prefiltered Baffin Bay seawater which was with F/2 medium and supplemented with antibiotics (0.1 g L^-1^ streptomycin, kanamycin, and antimycin; 0.05 g L^-1^ gentamicin). To evaluate the lower limits of phototrophy, triplicate cultures were acclimated for a minimum of 21 days to six constant irradiances (0.1, 0.3, 1, 3, 15, and 30 µmol photons m^-2^ s^-1^) which were provided by cool-white LEDs with spectral adjustments to mimic the under-ice light environment. Cultures were maintained in the exponential phase (∼1×10^6^ cells mL^-1^) by semi-continuous dilution to minimize biomass-induced light attenuation.

### Growth and Cellular Composition

Cell concentration and size were measured via a Beckman Coulter Multisizer 4 (Beckman Coulter, Brea, CA, USA). While cultures were maintained in exponential growth, the specific growth rates (μ, d^-1^) were calculated via a linearized exponential growth equation, with growth parameters derived by least-squares regression: lnN = ln(N_0_) + μT (Woods et al., 2005). The relationship between μ and irradiance was modelled to derive the maximum growth rate (μ_max_), the saturation parameter (𝐸^*K*^_μ_) and the compensation point for growth (I_c_) (MacIntyre et al., 2002). Cell viability was assessed via SYTOX Green stain (Sciandra et al., 2022). After acclimation, samples were harvested for the quantification of particulate organic carbon and nitrogen via elemental analysis (Costech ECS 8020), and pigment profiles via HPLC (Agilent 1260) following specific protocols (Guerin et al., 2022).

### Photosynthetic Performance

Photophysiology was assessed using variable chlorophyll *a* fluorescence and ^14^C-uptake assays. PSII activity was assessed with a Phyto-PAM fluorometer (Phyto-ML, Heinz Walz GmbH, Germany). Samples were dark-adapted for 30 min before exposure to Rapid Light Curves (RLC). Actinic light was provided by a modulated blue LED source (450 ± 20 nm) across incremental steps from 0.3 to 163 µmol photons m^-2^ s^-1^. The relative distribution of absorbed excitation energy in PSII was determined by the complementary PSII quantum yields method: ϕPSII = (F_m′_-F′)/F_m′_; ϕNPQ = F′/F_m′_ −F′/F_M_; and ϕNO = F′/F_M_ (Hendrickson et al., 2005; Klughammer and Schreiber, 2008; Xu et al., 2019). Additionally, the functional absorption cross-section of PSII (σPSII) and the maximum quantum yield of PSII (F_V_/F_M_) from cells after 30 min of dark acclimation were determined with a Fluorescence Induction and Relaxation fluorometer (FIRe, Satlantic, Canada). This instrument applies a saturating, single turnover flash (STF, 120μs) of blue light (455 nm) at saturation. The analysis of raw curves was performed within MATLAB software via the FIReWORX script by Audrey Barnett (https://sourceforge.net/projects/fireworx/). Carbon fixation rates were measured via ^14^C-bicarbonate incubations across 28 light levels (0-250 µmol photons m^-2^ s^-1^) with an in house incubator (Joli et al., 2024; Morin et al., 2020). A hyperbolic tangent model was fitted to the photosynthesis-irradiance data (Jassby & Platt, 1976) to derive the photosynthetic efficiency (α) and the maximum fixation rate (P_max_).

### Transcriptomics

Total RNA was extracted (Qiagen RNeasy) and DNAse-treated as described in Joli et al., 2024. Library preparation, which included poly-A selection, was performed using the Illumina TruSeq Stranded mRNA protocol. The generation of sequences was conducted and provided > 30 million paired-end reads (2 × 150 bp) per sample on an Illumina HiSeq 4000 platform. Read quality was verified using FastQC v0.11.9. Reads were aligned to the *F. cylindrus* reference genome (v1.0; Mock et al., 2017) using STAR v2.7.9a (Dobin et al., 2013). Raw read counts over exons were obtained with HTSeq-count v0.13.5 (Anders et al., 2015). Differential expression analysis was performed via DESeq2 v1.26.0 (Love et al., 2014) relative to the 30 µmol photons m^-2^ s^-1^ condition, with significance defined as |log2FC|>1 and an adjusted p-value < 0.05. The analysis targeted in-house manually annotated gene sets of 1 663 genes.

### Statistical Analysis

Data were analyzed using one-way ANOVA followed by a Tukey’s HSD post-hoc test, or Kruskal-Wallis followed by a Dunn’s test when assumptions of normality or homoscedasticity were not met. The significance threshold was defined at P < 0.05.

## ACKNOWLEDGMENTS

This work was supported by Laval University, CNRS/Sorbonne University, and the Agence Nationale de la Recherche (DIM: ANR-21-CE02-0021). We thank Québec Océan and Sentinel North (CFREF project) for their support. M.B. and C.B. thank the HFSP project, M.B. acknowledges an NSERC Discovery Grant, and C.B. thanks the Fondation BNP Paribas. Special thanks go to Dany Croteau, Benjamin Bailleul, Eric Marechal, and Doug Campbell for the rich discussions and their contributions to the analysis of these results. We also thank Juliette Laude and Petra Bulankova for useful discussions on the diatom cell cycle. We are grateful to Lena Bodiguel for her help with data analysis, fruitful discussion and code development, and to Carole-Anne Guay for her help with sample collection.

## REFERENCES

1. Agarwal A, Levitan O, Cruz De Carvalho H, Falkowski PG. 2023. Light-dependent signal transduction in the marine diatom Phaeodactylum tricornutum. Proceedings of the National Academy of Sciences 120: e2216286120.

2. Alou-Font E, Mundy C, Roy S, Gosselin M, Agustí S. 2013. Snow cover affects ice algal pigment composition in the coastal Arctic Ocean during spring. Marine Ecology Progress Series 474: 89–104.

3. Anders S, Pyl PT, Huber W. 2015. HTSeq—a Python framework to work with high-throughput sequencing data. Bioinformatics 31: 166–169.

4. Ardyna M, Arrigo KR. 2020. Phytoplankton dynamics in a changing Arctic Ocean. Nature Climate Change 10: 892–903.

5. Arrigo KR, Mills MM, Kropuenske LR, Van Dijken GL, Alderkamp A-C, Robinson DH. 2010. Photophysiology in Two Major Southern Ocean Phytoplankton Taxa: Photosynthesis and Growth of Phaeocystis antarctica and Fragilariopsis cylindrus under Different Irradiance Levels. Integrative and Comparative Biology 50: 950–966.

6. Aslam SN, Strauss J, Thomas DN, Mock T, Underwood GJC. 2018. Identifying metabolic pathways for production of extracellular polymeric substances by the diatom Fragilariopsis cylindrus inhabiting sea ice. The ISME Journal 12: 1237–1251.

7. Aslam SN, Underwood GJC, Kaartokallio H, Norman L, Autio R, Fischer M, Kuosa H, Dieckmann GS, Thomas DN. 2012. Dissolved extracellular polymeric substances (dEPS) dynamics and bacterial growth during sea ice formation in an ice tank study. Polar Biology 35: 661–676.

8. Bailleul B, Rogato A, De Martino A, Coesel S, Cardol P, Bowler C, Falciatore A, Finazzi G. 2010. An atypical member of the light-harvesting complex stress-related protein family modulates diatom responses to light. Proceedings of the National Academy of Sciences 107: 18214–18219.

9. Baldisserotto C, Ferroni L, Andreoli C, Fasulo MP, Bonora A, Pancaldi S. 2005. Dark-acclimation of the Chloroplast in Koliella antarctica Exposed to a Simulated Austral Night Condition. Arctic, Antarctic, and Alpine Research 37: 146–156.

10. Banse K. 2004. SHOULD WE CONTINUE TO USE THE 1% LIGHT DEPTH CONVENTION FOR ESTIMATING THE COMPENSATION DEPTH OF PHYTOPLANKTON FOR ANOTHER 70 YEARS? Limnology and Oceanography Bulletin 13: 49–52.

11. Beardall J, Morris I. 1976. The concept of light intensity adaptation in marine phytoplankton: Some experiments with Phaeodactylum tricornutum. Marine Biology 37: 377–387.

12. Behrenfeld MJ, Boss ES. 2018. Student’s tutorial on bloom hypotheses in the context of phytoplankton annual cycles. Global Change Biology 24: 55–77.

13. Bender ML, Zhu X-G, Falkowski P, Ma F, Griffin K. 2022. On the rate of phytoplankton respiration in the light. Plant Physiology 190: 267–279.

14. Castellani G, Veyssière G, Karcher M, Stroeve J, Banas SN, Bouman AH, Brierley SA, Connan S, Cottier F, Große F, et al. 2022. Shine a light: Under-ice light and its ecological implications in a changing Arctic Ocean. Ambio 51: 307–317.

15. Cohen JH, Berge J, Moline MA, Johnsen G, Zolich AP. 2020. Light in the Polar Night. In: Berge J, Johnsen G, Cohen JH, eds. Advances in Polar Ecology. POLAR NIGHT Marine Ecology. Cham: Springer International Publishing, 37–66.

16. Cota GF. 1985. Photoadaptation of high Arctic ice algae. Nature 315: 219–222.

17. Croteau D, Guérin S, Bruyant F, Ferland J, Campbell DA, Babin M, Lavaud J. 2021. Contrasting nonphotochemical quenching patterns under high light and darkness aligns with light niche occupancy in Arctic diatoms. Limnology and Oceanography 66.

18. Croteau D, Jaubert M, Falciatore A, Bailleul B. 2025. Pennate diatoms make non-photochemical quenching as simple as possible but not simpler. Nature Communications 16: 2385.

19. Croteau D, Jensen E, Wilhelm C, Bailleul B. 2024. Comparing Diatom Photosynthesis with the Green Lineage: Electron Transport, Carbon Fixation and Metabolism. In: Goessling JW, Serôdio J, Lavaud J, eds. Diatom Photosynthesis. Wiley, 1–44.

20. Dobin A, Davis CA, Schlesinger F, Drenkow J, Zaleski C, Jha S, Batut P, Chaisson M, Gingeras TR. 2013. STAR: ultrafast universal RNA-seq aligner. Bioinformatics 29: 15–21.

21. Dubinsky Z, Falkowski PG, Wyman K. 1986. Light Harvesting and Utilization by Phytoplankton. Plant and Cell Physiology 27: 1335–1349.

22. Dubinsky Z, Stambler N. 2009. Photoacclimation processes in phytoplankton: mechanisms, consequences, and applications. Aquatic Microbial Ecology 56: 163–176.

23. Falkowski PG, Owens TG. 1980. Light—Shade Adaptation: TWO STRATEGIES IN MARINE PHYTOPLANKTON. Plant Physiology 66: 592–595.

24. Fisher T, Shurtz-Swirski R, Gepstein S, Dubinsky Z. 1989. Changes in the Levels of Ribulose-l,5-bisphosphate Carboxylase/Oxygenase (Rubisco) in Tetraedron minimum (Chlorophyta) during Light and Shade Adaptation. Plant and Cell Physiology 30: 221–228.

25. Geider RJ, Osbonie BA, Raven JA. 1986. GROWTH, PHOTOSYNTHESIS AND MAINTENANCE METABOLIC COST IN THE DIATOM PHAEODACTYLUM TRICORNUTUM AT VERY LOW LIGHT LEVELS^1^. Journal of Phycology 22: 39–48.

26. Giossi CE, Kroth PG, Lepetit B. 2025a. Xanthophyll cycling and fucoxanthin biosynthesis in the model diatom Phaeodactylum tricornutum: recent advances and new gene functions. Frontiers in Photobiology 3: 1680034.

27. Giossi CE, Wünsch MA, Dautermann O, Schober AF, Buck JM, Kroth PG, Lohr M, Lepetit B. 2025b. Both major xanthophyll cycles present in nature promote nonphotochemical quenching in a model diatom. Plant Physiology 199: kiaf371.

28. Guérin S, Raguénès L, Croteau D, Babin M, Lavaud J. 2022. Potential for the Production of Carotenoids of Interest in the Polar Diatom Fragilariopsis cylindrus. Marine Drugs 20: 491.

29. Halsey KH, Jones BM. 2015. Phytoplankton Strategies for Photosynthetic Energy Allocation. Annual Review of Marine Science 7: 265–297.

30. Hancke K, Lund-Hansen LC, Lamare ML, Højlund Pedersen S, King MD, Andersen P, Sorrell BK. 2018. Extreme Low Light Requirement for Algae Growth Underneath Sea Ice: A Case Study From Station Nord, NE Greenland. Journal of Geophysical Research: Oceans 123: 985–1000.

31. Hellebust JA, Terborgh J. 1967. EFFECTS OF ENVIRONMENTAL CONDITIONS ON THE RATE OF PHOTOSYNTHESIS AND SOME PHOTOSYNTHETIC ENZYMES IN DUNALIELLA TERTIOLECTA BUTCHER1. Limnology and Oceanography 12: 559–567.

32. Hendrickson L, Förster B, Pogson BJ, Chow WS. 2005. A simple chlorophyll fluorescence parameter that correlates with the rate coefficient of photoinactivation of Photosystem II. Photosynthesis Research 84: 43–49.

33. Hoppe CJM, Fuchs N, Notz D, Anderson P, Assmy P, Berge J, Bratbak G, Guillou G, Kraberg A, Larsen A, et al. 2024. Photosynthetic light requirement near the theoretical minimum detected in Arctic microalgae. Nature Communications 15: 7385.

34. J. Geider R, Osborne BA, Raven JA. 1985. LIGHT DEPENDENCE OF GROWTH AND PHOTOSYNTHESIS IN PHAEODACTYLUM TRICORNUTUM (BACILLARIOPHYCEAE)^1^. Journal of Phycology 21: 609–619.

35. Jassby AD, Platt T. 1976. Mathematical formulation of the relationship between photosynthesis and light for phytoplankton. Limnology and Oceanography 21: 540–547.

36. Joli N, Concia L, Mocaer K, Guterman J, Laude J, Guerin S, Sciandra T, Bruyant F, Ait-Mohamed O, Beguin M, et al. 2024. Hypometabolism to survive the long polar night and subsequent successful return to light in the diatom Fragilariopsis cylindrus. New Phytologist 241: 2193–2208.

37. Juchem DP, Schimani K, Holzinger A, Permann C, Abarca N, Skibbe O, Zimmermann J, Graeve M, Karsten U. 2023. Lipid degradation and photosynthetic traits after prolonged darkness in four Antarctic benthic diatoms, including the newly described species Planothidium wetzelii sp. nov. Frontiers in Microbiology 14: 1241826.

38. Kennedy F, Martin A, Bowman JP, Wilson R, McMinn A. 2019. Dark metabolism: a molecular insight into how the Antarctic sea-ice diatom Fragilariopsis cylindrus survives long-term darkness. New Phytologist 223: 675–691.

39. Khuller AR, Warren SG, Christensen PR, Clow GD. 2024. Potential for photosynthesis on Mars within snow and ice. Communications Earth & Environment 5: 583.

40. Kolber ZS, Prášil O, Falkowski PG. 1998. Measurements of variable chlorophyll fluorescence using fast repetition rate techniques: defining methodology and experimental protocols. Biochimica et Biophysica Acta (BBA) - Bioenergetics 1367: 88–106.

41. Kristiansen T, Varpe Ø, Selig ER, Laurel BJ, Sydeman WJ, Hegglin MI, Wallhead PJ. 2025. Climate change impacts on ocean light in Arctic ecosystems. Nature Communications 16: 9798.

42. Kropuenske LR, Mills MM, Van Dijken GL, Bailey S, Robinson DH, Welschmeyer NA, Arrigoa KR. 2009. Photophysiology in two major Southern Ocean phytoplankton taxa: Photoprotection in Phaeocystis antarctica and Fragilariopsis cylindrus. Limnology and Oceanography 54: 1176–1196.

43. Kughammer C, Schreiber U. 2008. Complementary PS II quantum yields calculated from simple fluorescence parameters measured by PAM fluorometry and the saturation pulse method. PAM application notes: 27–35.

44. Kvernvik AC, Hoppe CJM, Lawrenz E, Prášil O, Greenacre M, Wiktor JM, Leu E. 2018. Fast reactivation of photosynthesis in arctic phytoplankton during the polar night^1^ (K Valentin, Ed.). Journal of Phycology 54: 461–470.

45. Lacour T, Babin M, Lavaud J. 2020. Diversity in Xanthophyll Cycle Pigments Content and Related Nonphotochemical Quenching (NPQ) Among Microalgae: Implications for Growth Strategy and Ecology (P Kroth, Ed.). Journal of Phycology 56: 245–263.

46. Lacour T, Larivière J, Babin M. 2017. Growth, Chl a content, photosynthesis, and elemental composition in polar and temperate microalgae. Limnology and Oceanography 62: 43–58.

47. Lacour T, Larivière J, Ferland J, Morin P-I, Grondin P-L, Donaher N, Cockshutt A, Campbell DA, Babin M. 2022. Photoacclimation of the polar diatom Chaetoceros neogracilis at low temperature (A Quigg, Ed.). PLOS ONE 17: e0272822.

48. Lafond A, Leblanc K, Quéguiner B, Moriceau B, Leynaert A, Cornet V, Legras J, Ras J, Parenteau M, Garcia N, et al. 2019. Late spring bloom development of pelagic diatoms in Baffin Bay (JW Deming and C Michel, Eds). Elementa: Science of the Anthropocene 7: 44.

49. Langdon C. 1987. On the causes of interspecific differences in the growth-irradiance relationship for phytoplankton. Part I. A comparative study of the growth-irradiance relationship of three marine phytoplankton species: Skeletonema costatum, Olisthodiscus luteus and Gonyaulax tamarensis. Journal of Plankton Research 9: 459–482.

50. Lavaud J, Goss R. 2014. The Peculiar Features of Non-Photochemical Fluorescence Quenching in Diatoms and Brown Algae. In: Demmig-Adams B, Garab G, Adams Iii W, Govindjee, eds. Advances in Photosynthesis and Respiration. Non-Photochemical Quenching and Energy Dissipation in Plants, Algae and Cyanobacteria. Dordrecht: Springer Netherlands, 421–443.

51. Lavaud J, Rousseau B, Etienne A-L. 2002. In diatoms, a transthylakoid proton gradient alone is not sufficient to induce a non-photochemical fluorescence quenching. FEBS Letters 523: 163–166.

52. Laws EA, Bannister TT. 1980. Nutrient- and light-limited growth of Thalassiosira fluviatilis in continuous culture, with implications for phytoplankton growth in the ocean1. Limnology and Oceanography 25: 457–473.

53. Leu E, Mundy CJ, Assmy P, Campbell K, Gabrielsen TM, Gosselin M, Juul-Pedersen T, Gradinger R. 2015. Arctic spring awakening – Steering principles behind the phenology of vernal ice algal blooms. Progress in Oceanography 139: 151–170.

54. Lewis KM, Van Dijken GL, Arrigo KR. 2020. Changes in phytoplankton concentration now drive increased Arctic Ocean primary production. Science 369: 198–202.

55. Lovejoy C, Legendre L, Martineau M-J, Bâcle J, Von Quillfeldt CH. 2002. Distribution of phytoplankton and other protists in the North Water. Deep Sea Research Part II: Topical Studies in Oceanography 49: 5027–5047.

56. MacIntyre HL, Kana TM, Anning T, Geider RJ. 2002. PHOTOACCLIMATION OF PHOTOSYNTHESIS IRRADIANCE RESPONSE CURVES AND PHOTOSYNTHETIC PIGMENTS IN MICROALGAE AND CYANOBACTERIA^1^. Journal of Phycology 38: 17–38.

57. Marra JF, Chamberlin WS, Knudson CA, Rhea WJ, Ho C. 2023. Parameters for the depth of the ocean’s productive layer. Frontiers in Marine Science 10: 1052307.

58. Massicotte P, Amiraux R, Amyot M-P, Archambault P, Ardyna M, Arnaud L, Artigue L, Aubry C, Ayotte P, Bécu G, et al. 2020. Green Edge ice camp campaigns: understanding the processes controlling the under-ice Arctic phytoplankton spring bloom. Earth System Science Data 12: 151–176.

59. Michelle Wood A, Everroad RC, Wingard LM. 2005. Measuring Growth Rates in Microalgal Cultures. In: Algal Culturing Techniques. Elsevier, 269–285.

60. Mock T, Gradinger R. 1999. Determination of Arctic ice algal production with a new in situ incubation technique. Marine Ecology Progress Series 177: 15–26.

61. Mock T, Otillar RP, Strauss J, McMullan M, Paajanen P, Schmutz J, Salamov A, Sanges R, Toseland A, Ward BJ, et al. 2017. Evolutionary genomics of the cold-adapted diatom Fragilariopsis cylindrus. Nature 541: 536–540.

62. Morin P, Lacour T, Grondin P, Bruyant F, Ferland J, Forget M, Massicotte P, Donaher N, Campbell DA, Lavaud J, et al. 2020. Response of the sea-ice diatom Fragilariopsis cylindrus to simulated polar night darkness and return to light. Limnology and Oceanography 65: 1041–1060.

63. Palmisano AC, SooHoo JB, White DC, Smith GA, Stanton GR, Burckle LH. 1985. SHADE ADAPTED BENTHIC DIATOMS BENEATH ANTARCTIC SEA ICE^1^. Journal of Phycology 21: 664–667.

64. Poulin M, Daugbjerg N, Gradinger R, Ilyash L, Ratkova T, Von Quillfeldt C. 2011. The pan-Arctic biodiversity of marine pelagic and sea-ice unicellular eukaryotes: a first-attempt assessment. Marine Biodiversity 41: 13–28.

65. Quigg A, Beardall J. 2003. Protein turnover in relation to maintenance metabolism at low photon flux in two marine microalgae. Plant, Cell & Environment 26: 693–703.

66. Quigg A, Kevekordes K, Raven JA, Beardall J. 2006. Limitations on microalgal growth at very low photon fluence rates: the role of energy slippage. Photosynthesis Research 88: 299–310.

67. Randelhoff A, Lacour L, Marec C, Leymarie E, Lagunas J, Xing X, Darnis G, Penkerc’h C, Sampei M, Fortier L, et al. 2020. Arctic mid-winter phytoplankton growth revealed by autonomous profilers. Science Advances 6: eabc2678.

68. Raven JA, Beardall J. 2016. Dark Respiration and Organic Carbon Loss. In: Borowitzka MA, Beardall J, Raven JA, eds. The Physiology of Microalgae. Cham: Springer International Publishing, 129–140.

69. Raven JA, Kübler JE, Beardall J. 2000. Put out the light, and then put out the light. Journal of the Marine Biological Association of the United Kingdom 80: 1–25.

70. Richardson K, Fogg GE. 1982. The role of dissolved organic material in the nutrition and survival of marine dinoflagellates. Phycologia 21: 17–26.

71. Schaub I, Wagner H, Graeve M, Karsten U. 2017. Effects of prolonged darkness and temperature on the lipid metabolism in the benthic diatom Navicula perminuta from the Arctic Adventfjorden, Svalbard. Polar Biology 40: 1425–1439.

72. Sciandra T, Forget M, Bruyant F, Béguin M, Lacour T, Bowler C, Babin M. 2022. The possible fates of Fragilariopsis cylindrus (polar diatom) cells exposed to prolonged darkness (J Raven, Ed.). Journal of Phycology 58: 281–296.

73. Suggett DJ, Le Floc’H E, Harris GN, Leonardos N, Geider RJ. 2007. Different strategies of photoacclimation by two strains of Emiliania huxleyi (Haptophyta)^1^. Journal of Phycology 43: 1209–1222.

74. Suzuki Y, Kudoh S, Takahashi M. 1997. Photosynthetic and respiratory characteristics of an Arctic ice algal community living in low light and low temperature conditions. Journal of Marine Systems 11: 111–121.

75. Sverdrup HU. 1953. On Conditions for the Vernal Blooming of Phytoplankton. ICES Journal of Marine Science 18: 287–295.

76. Ugalde SC, Meiners KM, Davidson AT, Westwood KJ, McMinn A. 2013. Photosynthetic carbon allocation of an Antarctic sea ice diatom (Fragilariopsis cylindrus). Journal of Experimental Marine Biology and Ecology 446: 228–235.

77. Underwood G, Fietz S, Papadimitriou S, Thomas D, Dieckmann G. 2010. Distribution and composition of dissolved extracellular polymeric substances (EPS) in Antarctic sea ice. Marine Ecology Progress Series 404: 1–19.

78. Van De Poll WH, Abi Nassif T. 2023. The interacting effect of prolonged darkness and temperature on photophysiological characteristics of three Antarctic phytoplankton species. Journal of Phycology 59: 1053–1063.

79. Xu K, Lavaud J, Perkins R, Austen E, Bonnanfant M, Campbell DA. 2018. Phytoplankton σPSII and Excitation Dissipation; Implications for Estimates of Primary Productivity. Frontiers in Marine Science 5: 281.

