## Supplemental Figures 1 to 10 for "Paradoxical energetics in the polar diatom *Fragilariopsis cylindrus* exposed to extreme low light"

**from**

### Figures

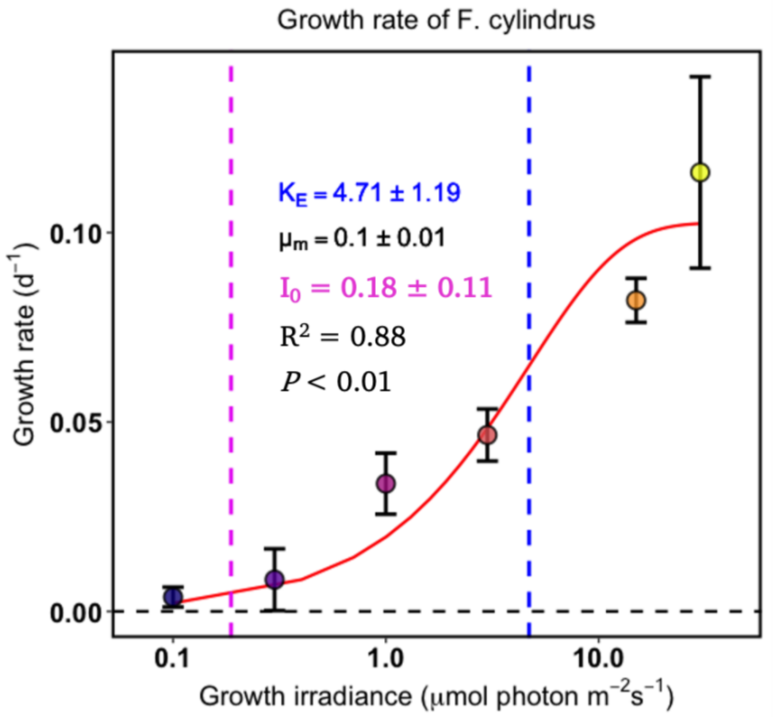


**Figure S1. Growth rate (μ, $d^{-1}$) as function of light intensity curve (μmol photon $m^{-2}$ $s^{-1}$) for the polar diatom *F. cylindrus*.** The dots represent the observed specific growth rates (μ) derived by the linearized exponential growth equation (Woods et al., 2005) and the red line shows the modelled relationship between the growth rate μ and E following (MacIntyre et al., 2002). $\mu_m$ is the maximum growth rate ($d^{-1}$). $K_E$ is the light saturation parameter for growth (μmol photons $m^{-2}$ $s^{-1}$) and $I_0$ the light compensation point for growth. derived by solving the equation with the minimum growth rates. The data representing the mean value and the error bar the standard deviation (n = 3).

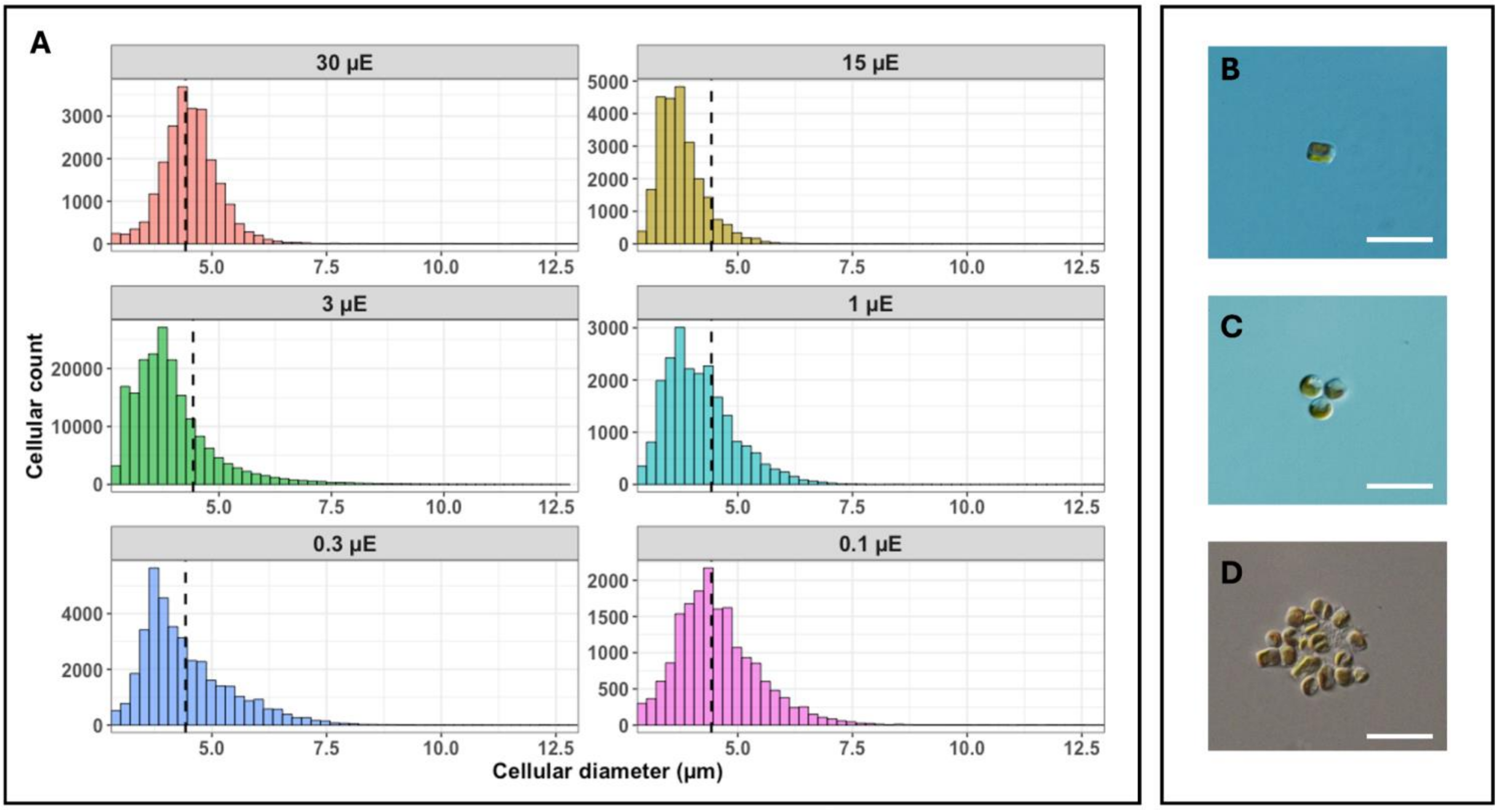


**Figure S2. Cell size distribution and aggregation phenotypes of *F. cylindrus* across light conditions**. (A) Cellular size distribution for all acclimated cells at the different light intensity (µE = µmol photon $m^{-2}$ $s^{-1}$). The dashed line represents the mean cellular size observed at 30 µmol photon $m^{-2}$ $s^{-1}$. (B-D) Microscopy images illustrating cellular phenotypes: (B) single cells observed at 30 µmol photon $m^{-2}$ $s^{-1}$, and (C-D) aggregated cells observed at the lowest light condition (0.1 µE). Scale bars = 10 µm.

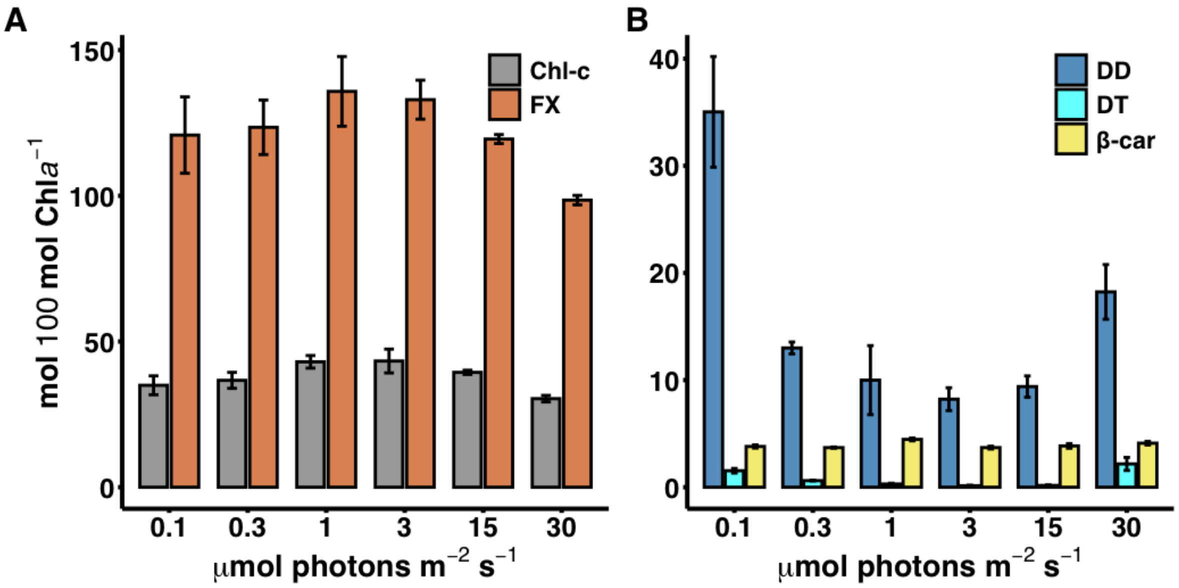


**Figure S3. Molar ratios of pigment normalized to 100 mol of chlorophyll *a* in cells acclimated to different growth irradiances.** (A) Photosynthetic accessory pigments, fucoxanthin (FX) and total chlorophyll *c* (Chl *c*, sum of $c_1$ and $c_2$). (B) Photoprotective pigments, diadinoxanthin (DD), diatoxanthin (DT), and β-carotene (β-car). Data represent the mean ± SD (n = 3).

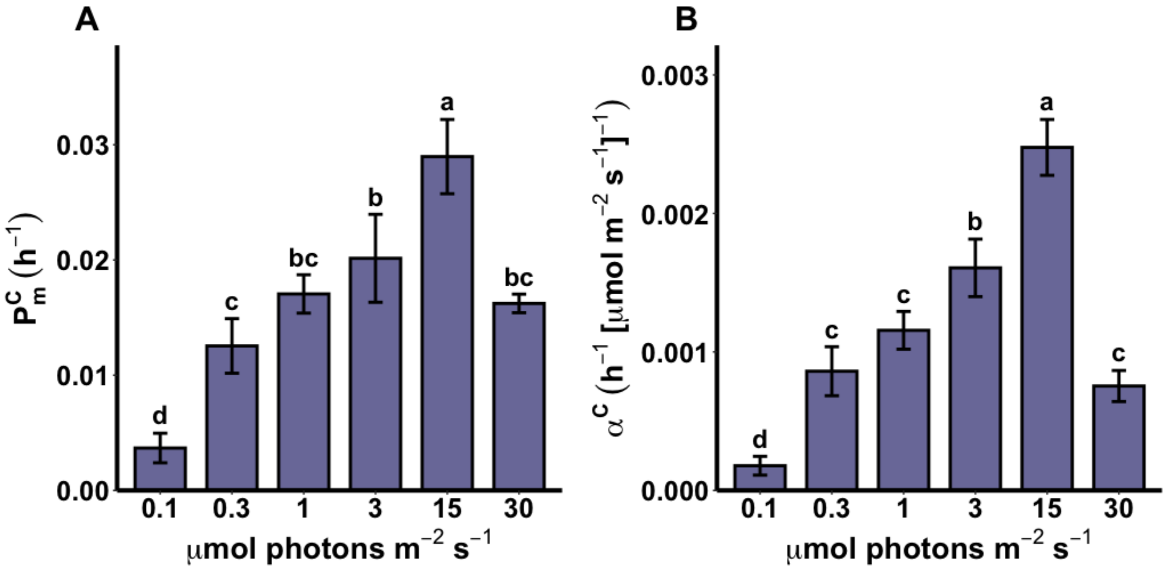


**Figure S4. Photosynthetic parameters derived from modeled carbon-specific photosynthesis–irradiance (PE) curves**. (A) Light-saturated carbon-specific photosynthetic rate ($P^{C}_{m}$ , in $h^{-1}$). (B) Carbon-specific initial slope of the P-E curve ($\alpha^C$ , in $h^{-1}$ [μ mol photon $m^{-2}$ $s^{-1}$]$^{-1}$). Parameters were estimated from $^{14}C$-bicarbonate uptake rates measured during 20-min incubations at each light level of the curve. Blue bars represent means (n = 3). Error bars represent standard deviation (SD), and different letters indicate significant differences between growth irradiances ($P<0.05$) determined by Tukey's HSD post hoc analysis.

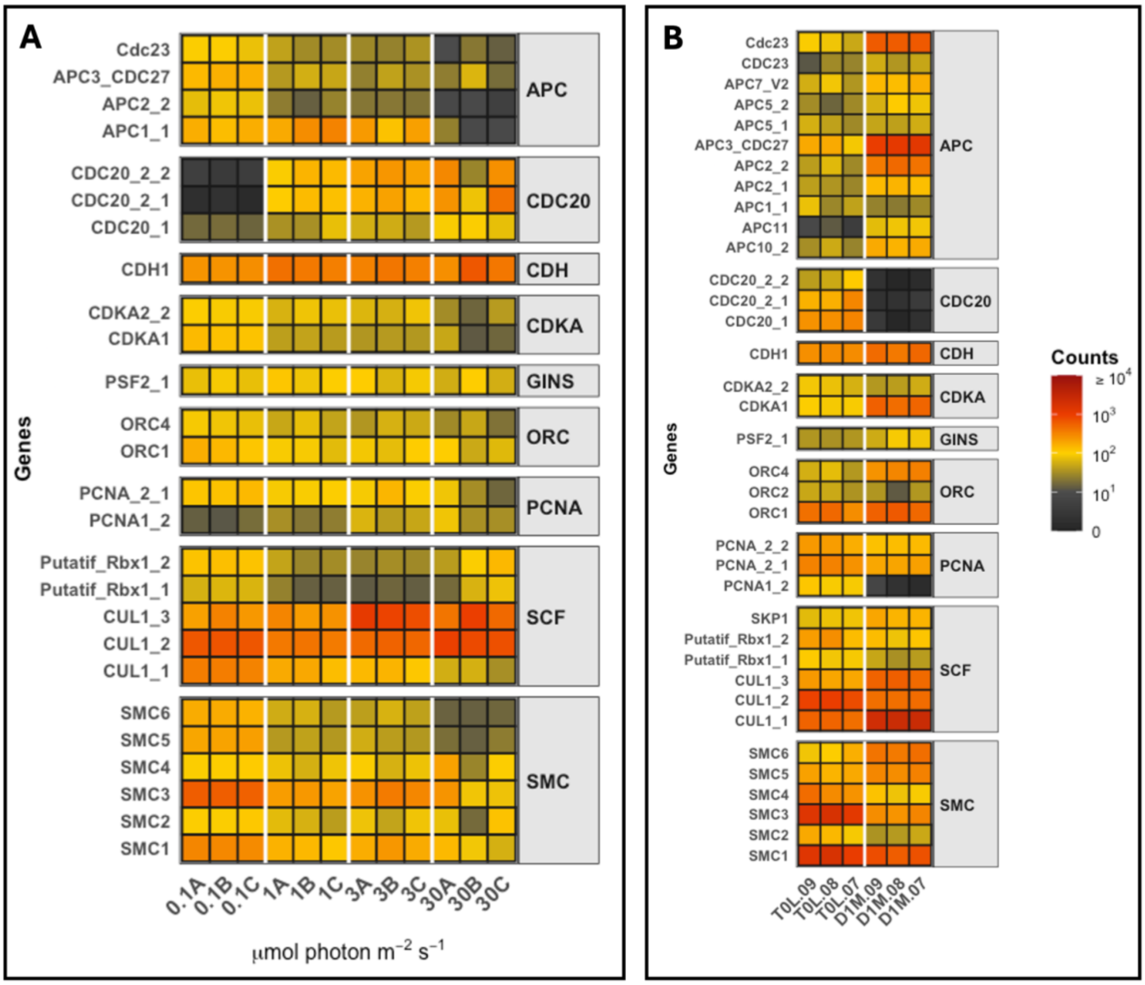


**Figure S5. Transcriptional patterns of cell cycle genes.** The heatmaps show the expression profiles of actively transcribed genes involved in the cell cycle (>30 DESeq2-normalized counts). (A) Transcriptional patterns from the current study after the acclimation period specific to each growth irradiance. (B) Comparative transcriptional patterns retrieved from Joli et al. (2024) for *F. cylindrus* (same strain) acclimated to 30 µmol photons $m^{-2}$ $s^{-1}$ (T0L) and after one month of complete darkness (D1M). The color gradient represents the expression level (DESeq2-normalized counts) for each triplicate per condition. Abbreviations: APC, anaphase-promoting complex; PCNA, proliferating cell nuclear antigen; ORC, origin recognition complex; SCF, Skp, Cullin, F-box containing complex. See dataset S1 for the corresponding JGI IG gene.

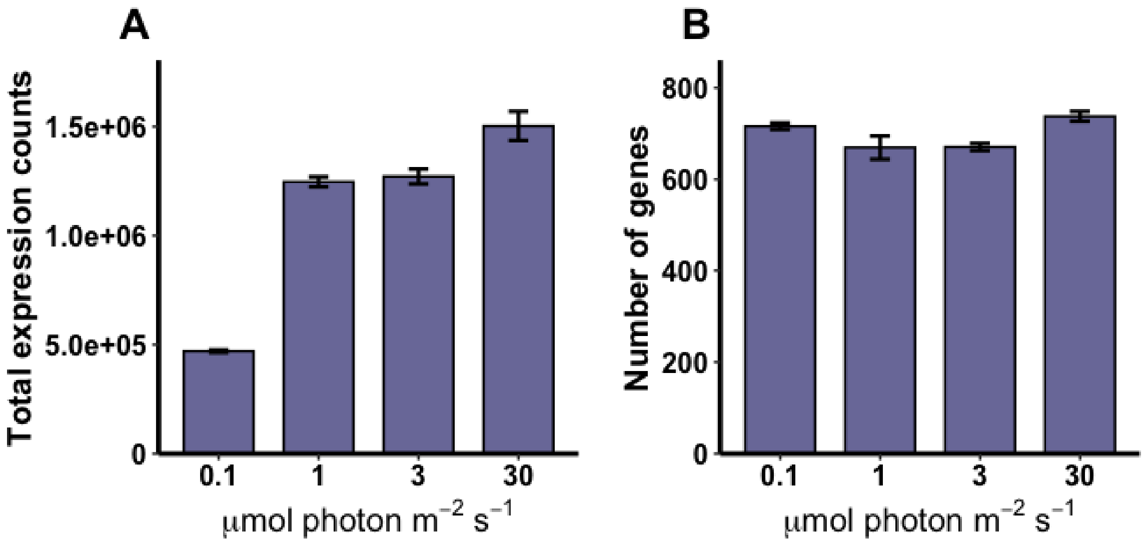


**Figure S6. Global transcriptional activity after the acclimation period specific to each growth irradiance.** (A) Number of actively transcribed genes per light condition. Genes were considered actively transcribed if their expression exceeded 30 DESeq2-normalized counts. (B) Total sum of normalized transcript counts per condition. Data represent the mean ± SD (n = 3).

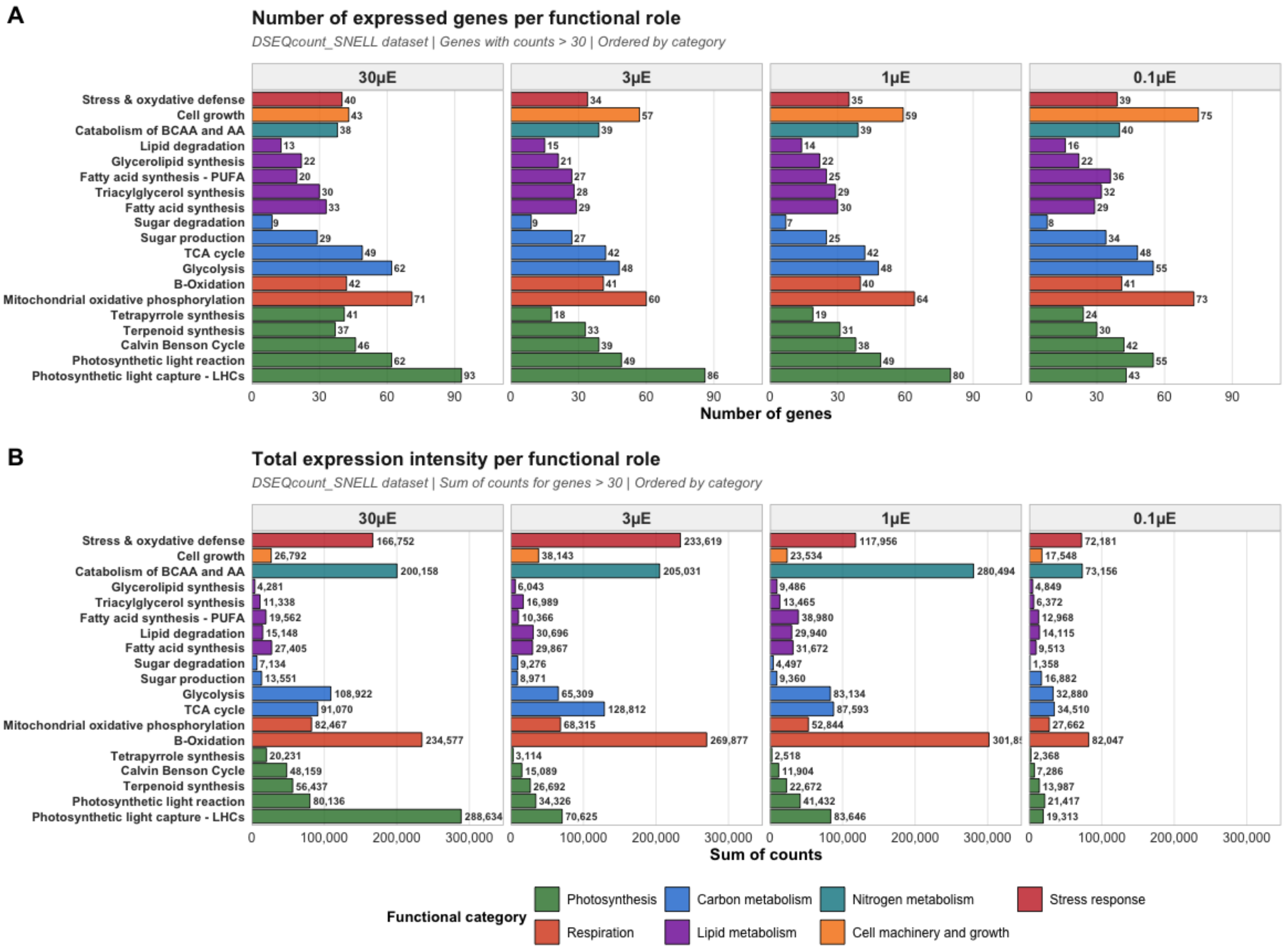


**Figure S7. Partitioning of transcriptional activity by broad metabolic functional categories after the acclimation period specific to each growth irradiance.** (A) Number of actively transcribed genes (>30 DESeq2-normalized counts) attributed to each major functional category across growth irradiances. (B) Cumulative abundance of DESeq2-normalized transcript assigned to each metabolic category. Functional groups correspond to the categories defined in Figure 4.

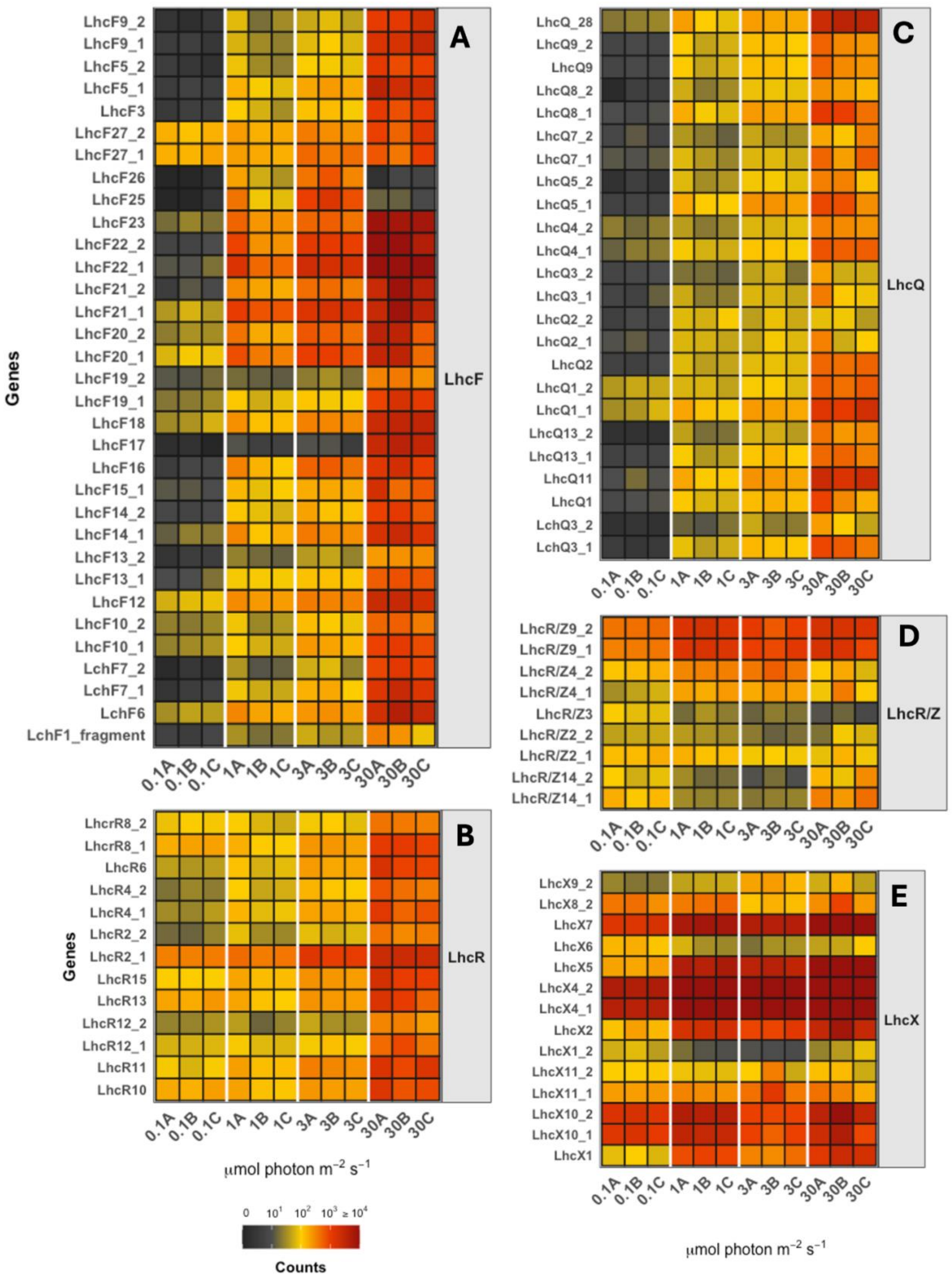
A
B
C
D
E
LhcF
LhcR
LhcQ
LhcR/Z
LhcX
Genes
LhcF9_2
LhcF9_1
LhcF5_2
LhcF5_1
LhcF3
LhcF27_2
LhcF27_1
LhcF26
LhcF25
LhcF23
LhcF22_2
LhcF22_1
LhcF21_2
LhcF21_1
LhcF20_2
LhcF20_1
LhcF19_2
LhcF19_1
LhcF18
LhcF17
LhcF16
LhcF15_1
LhcF14_2
LhcF14_1
LhcF13_2
LhcF13_1
LhcF12
LhcF10_2
LhcF10_1
LchF7_2
LchF7_1
LchF6
LchF1_fragment
LhcrR8_2
LhcrR8_1
LhcR6
LhcR4_2
LhcR4_1
LhcR2_2
LhcR2_1
LhcR15
LhcR13
LhcR12_2
LhcR12_1
LhcR11
LhcR10
LhcQ_28
LhcQ9_2
LhcQ9
LhcQ8_2
LhcQ8_1
LhcQ7_2
LhcQ7_1
LhcQ5_2
LhcQ5_1
LhcQ4_2
LhcQ4_1
LhcQ3_2
LhcQ3_1
LhcQ2_2
LhcQ2_1
LhcQ2
LhcQ1_2
LhcQ1_1
LhcQ13_2
LhcQ13_1
LhcQ11
LhcQ1
LchQ3_2
LchQ3_1
LhcR/Z9_2
LhcR/Z9_1
LhcR/Z4_2
LhcR/Z4_1
LhcR/Z3
LhcR/Z2_2
LhcR/Z2_1
LhcR/Z14_2
LhcR/Z14_1
LhcX9_2
LhcX8_2
LhcX7
LhcX6
LhcX5
LhcX4_2
LhcX4_1
LhcX2
LhcX1_2
LhcX11_2
LhcX11_1
LhcX10_2
LhcX10_1
LhcX1
0.1A 0.1B 0.1C 1A 1B 1C 3A 3B 3C 30A 30B 30C
μmol photon m⁻² s⁻¹
0 10¹ 10² 10³ ≥ 10⁴
Counts

**Figure S8. Transcriptional patterns of light-harvesting complex (LHC) genes**. The show display the expression profiles of actively transcribed genes (>30 DESeq2-normalized counts) encoding antenna protein fucoxanthin-chlorophyll a/c binding proteins (FCPs) in *F. cylindrus* after the acclimation period specific to each growth irradiance. The panels represent the expression values for distinct FCP families: (A) *lhcf* proteins representing the major PSII-related antenna complexes; (B) *lhcr* proteins associated with PSI; (C) *lhcq*; (D) *lhcz*; and (E) *lhcx* proteins responsible for energy-dependent non-photochemical quenching (NPQ). The color gradient represents the expression level (DESeq2-normalized counts) for each gene per biological triplicate and for each light condition. Corresponding JGI gene IDs are listed in Dataset S1.

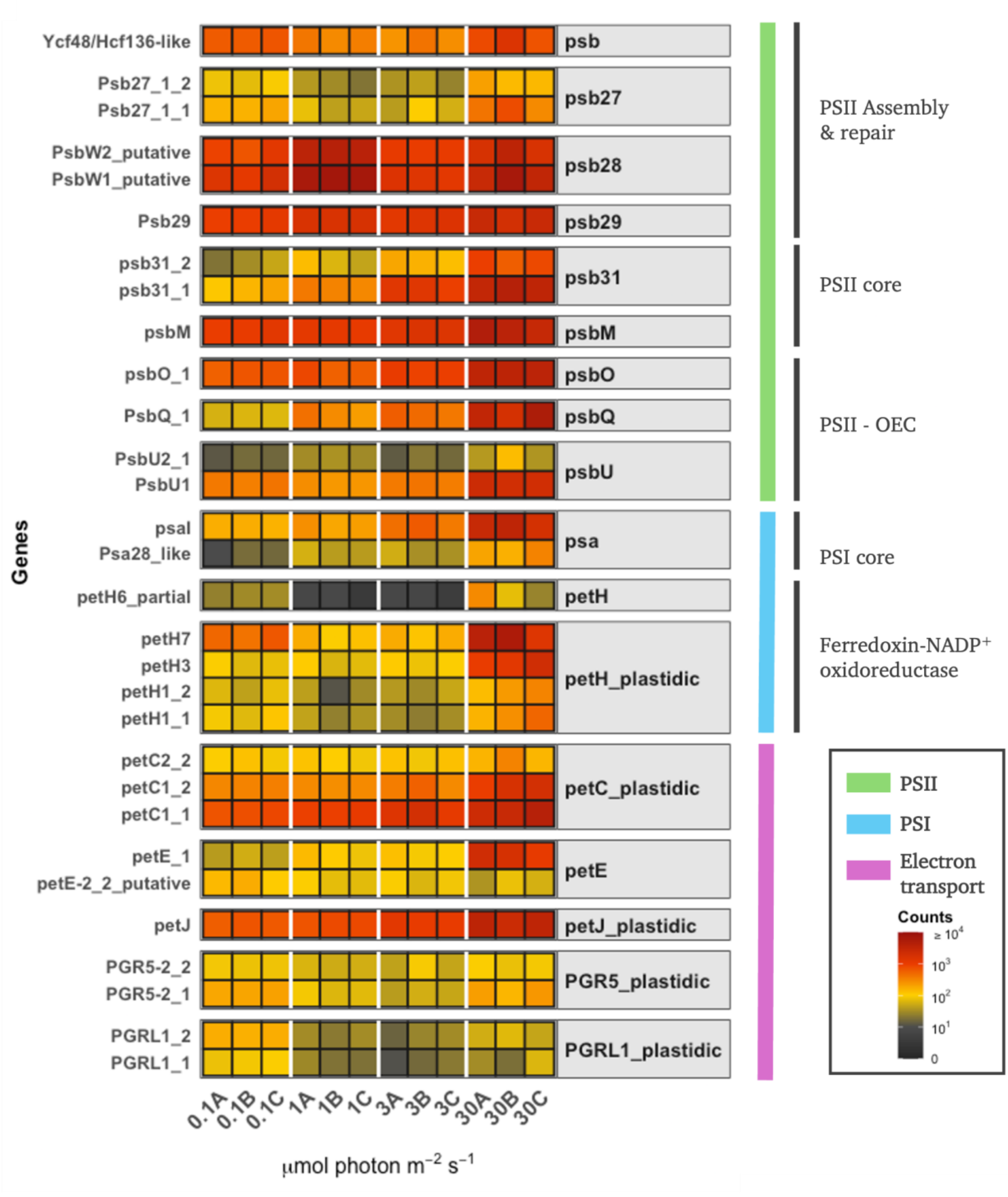


**Figure S9. Transcriptional patterns of photosynthetic light reaction genes.** The heatmaps display the expression profiles of actively transcribed genes (>30 DESeq2-normalized counts) encoding components of Photosystem II (PSII), Photosystem I (PSI), and the electron transport chain after the acclimation period specific to each growth irradiance. The color gradient represents the expression level (DESeq2-normalized counts) for each gene per biological triplicate and for each light condition. The color bar on the left side represents the corresponding affiliation of genes between the major components. Corresponding JGI gene IDs are listed in Dataset S1. Abbreviations: OEC, oxygen-evolving complex; petC, Rieske FeS center of the cytochrome $b_6f$ complex; petE, Plastocyanin; petJ, Cytochrome $c_6$; PGR5, proton gradient regulation 5; PGRL1, PGR5-like photosynthetic phenotype 1; psa, Photosystem I genes; psb, Photosystem II genes. Corresponding JGI gene IDs are listed in Dataset S1.

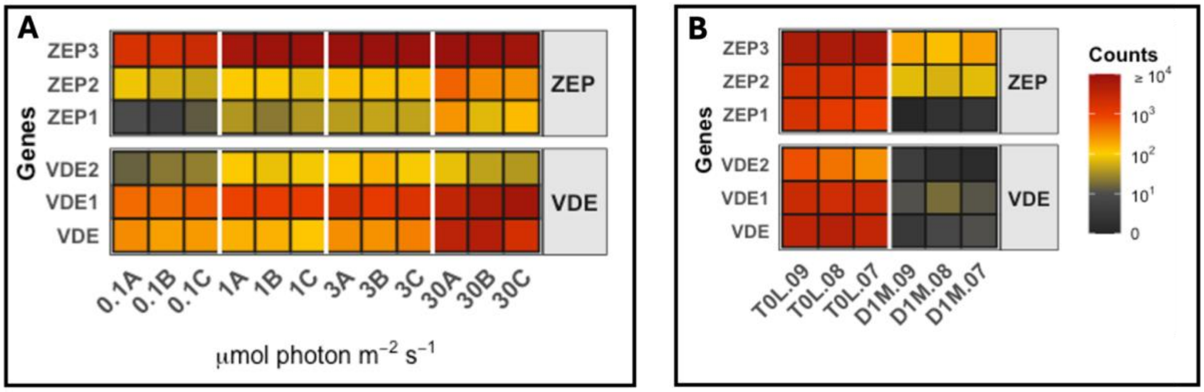


**Figure S10. Transcriptional patterns of xanthophyll cycle enzymes.** The heatmaps show the expression profiles of actively transcribed genes (>30 DESeq2-normalized counts) encoding the Violaxanthin De-Epoxidase (VDE) and Zeaxanthin Epoxidase (ZEP). (A) Expression profiles in F. cylindrus from the current study after the acclimation period specific to each growth irradiance. (B) Comparative transcriptional patterns retrieved from Joli et al. (2024) for *F. cylindrus* (same strain) acclimated to 30 µmol photons $m^{-2}$ $s^{-1}$ (T0L) and after one month of complete darkness (D1M). The color gradient represents the expression level (DESeq2-normalized counts) for each gene per biological triplicate and for each light condition. Corresponding JGI gene IDs are listed in Dataset S1.
